# Nanoscale dendritic shaft constrictions shape synaptic integration in fine caliber principal neuron dendrites

**DOI:** 10.1101/2025.07.18.664460

**Authors:** Tony Kelly, Michael Döngi, Juan Eduardo Rodriguez-Gatica, Netanel Ofer, Carlos Wert-Carvajal, Michela Barboni, Philipp Bethge, Michel Herde, Jens Tillmann, Sabrina Ingrid Peter, Sebastian Dupraz, Henner Koch, Valentin Stein, Frank Bradke, Tatjana Tchumatchenko, Martin Karl Schwarz, Ulrich Kubitscheck, U. Valentin Nägerl, Heinz Beck

## Abstract

Traditionally, theoretical studies typically described dendritic morphology as optimized for efficient synaptic voltage transfer from spines to the soma, implemented as a tubular design respecting Rall’s 3/2 rule for impedance matching at branch points. Here, we reveal that this view is an oversimplification. Using three high-resolution imaging techniques, we demonstrate that dendrites in cortical and hippocampal neurons contain nanoscale constrictions, comparable in diameter to spine necks. We provide theoretical and experimental evidence that these constrictions partition the dendrite into distinct electrical compartments, significantly shaping dendritic integration of synaptic potentials.

## Main Text

Dendrites receive thousands of excitatory inputs directed to dendritic spines. First described by Ramón y Cajal in 1888, dendritic spines exhibit a distinctive nano-anatomical structure, consisting of a bulbous head connected to the dendrite by an elongated spine neck. Spine necks, with diameters ranging from 90 to several hundred nanometers, create electrochemical compartments essential for synaptic function and plasticity (reviewed in ^1–5^). In contrast, dendrites have traditionally been considered as continuous tubes lacking such submicron specializations along their axial length ^6,7^. Here, we challenge this view by presenting high-resolution imaging evidence for nanoscale dendritic shaft constrictions (DSCs). Using computational modeling and two-photon glutamate uncaging, we also show that these newly identified structures have a significant impact on EPSPs in the dendrite and soma.

First, we used expansion microscopy (ExM) in conjunction with a retroviral approach to sparsely label hippocampal dentate granule cells (**Fig. 1a**), and reconstructed entire individual dendrites from the soma to their apical tips (n=6 dendrites, n=5 cells and n=4 animals, **Fig. 1a, Extended Data Fig. 1a, b**). The average dendritic diameters decreased from proximal (0.80±0.07 µm at 1-100 µm from the soma) to distal dendritic segments (0.62±0.08 µm at 100-200 µm, 0.54±0.07 µm at >200 µm, **Extended Data Fig. 1b, d**). However, we also observed local reductions in dendritic diameter within individual branches where the dendrite would be markedly thinner than in the adjacent segments (**Fig. 1a, right panels, Extended Data Fig. 1b, c**). The local reductions were defined as abrupt drops in dendritic diameter by twofold or more which then recovered to at least 75% of the initial dendritic diameter. We termed these structures dendritic shaft constrictions (DSCs). DSCs on average were found to have a z-scored diameter change of -1.97±0.1 (n=35 DSCs in 6 dendrites, see **Extended Data Fig. 1b**). The average DSC diameter was 0.33±0.03 µm (range 0.11-0.46 µm, n=35 DSCs in 6 dendrites, **Fig. 1d**), while their average length was 1.4±0.2 µm (distribution see **Fig. 1e**). Most DSCs were observed distally, with the most proximal DSC being on average 207±18 µm (range 56-249 µm, n=6) away from the soma (**Fig. 1f, Extended Data Fig. 1b**). We counted on average 5 (range 2-10) DSCs per individual GC dendrite, which, when summed up per dendrite corresponded to a total DSC length of 8.6±2.6 µm/dendrite (range 3.4-21 µm). Multiple DSCs created inter-DSC compartments, which were on average 17±1.5 µm long (distribution see **Fig. 1g**). Based on basic biophysical principles, these morphological constrictions are expected to increase the axial resistance of the dendrite by more than 5-fold from 5.5±1.9 MΩ to 34.8±6.0 MΩ per µm length of dendrite (n=6 dendrites and 35 DSCs; ^8^) adding a total of 260±114 MΩ in axial resistance (range 58–771 MΩ) to an individual dendrite (n=6 dendrites).

**Fig. 1:**
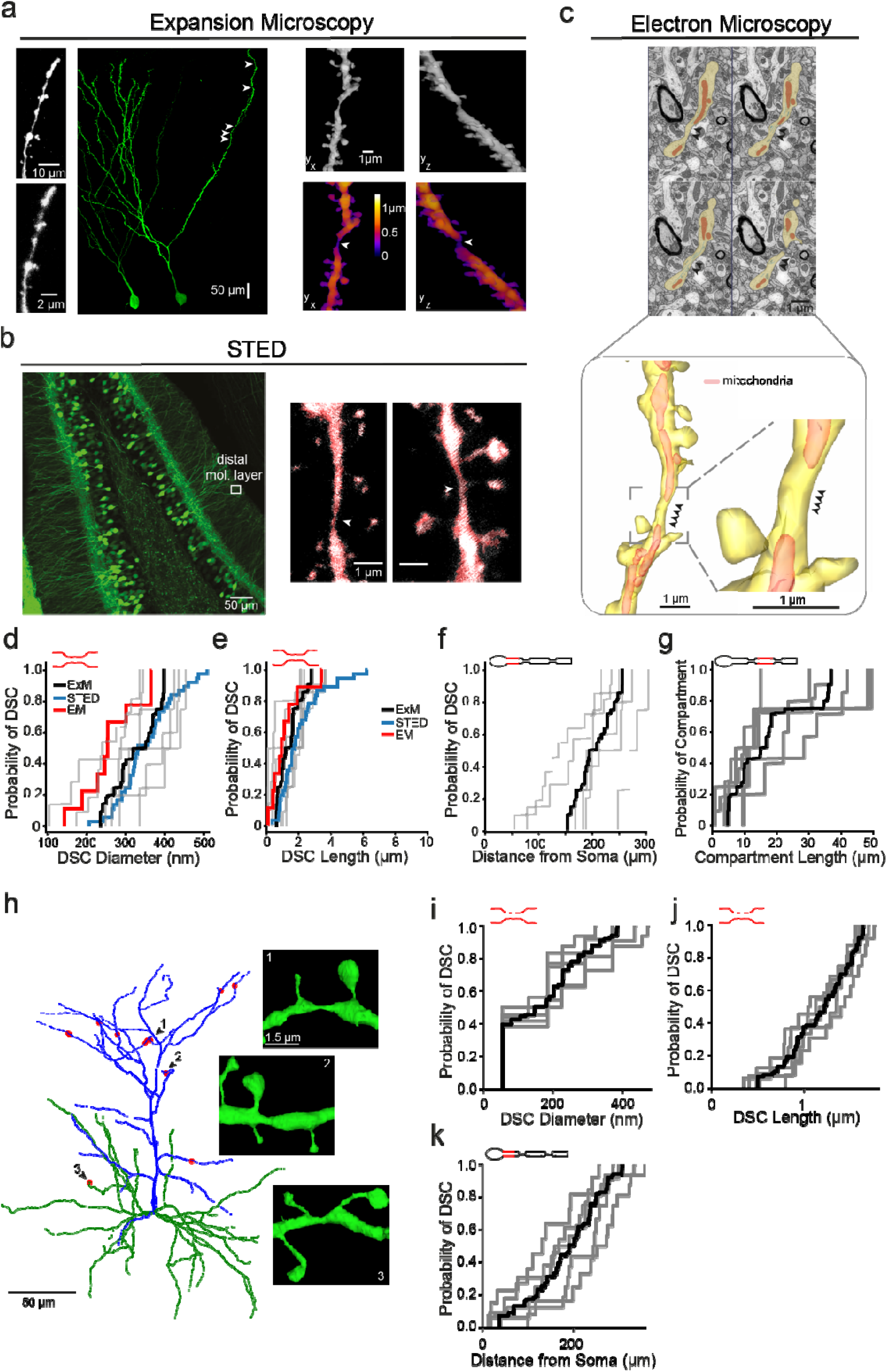
Dendritic nanoscale shaft constrictions (DSCs) are a common feature of hippocampal and cortical dendrites. **a**, Expansion microcopy (ExM). Left: Light-sheet fluorescence microscopy (LSFM) maximum projection image of granule cells sparsely labelled with eGFP. Using expansion microscopy (ExM) tissue was expanded by 3.8 times to virtually improve optical resolution. *Inserts* at far left show enlargements of same dendritic segment pre- and post-expansion. Locations of identified dendritic shaft constriction (DSCs) on reconstructed dendrite indicated with arrows. Right: Two examples of 3D reconstructions as grey scale image (*above*) or pseudo-colored with the diameter of the largest non-overlapping sphere (*below, see Methods*). DSCs indicated with arrows. Scale bar is corrected for expansion factor. b, STED imaging. Left: Confocal maximum projection image of dentate gyrus from a Thy1-YFP expressing mouse. Right: STED maximum projection grey scale images with overlapping binary image (red) of short dendritic segments from the distal molecular layer indicated in left panel. DSCs indicated with arrows. **c**, Electron microscopy (EM). Top: Series of 4 electron micrographs from a stack taken from the outer molecular layer (172 µm from the granule cell layer). Segmented dendrite in yellow, DSC indicated with black arrowheads, mitochondria labelled in red. Bottom: 3D reconstruction of the dendrite and enlargement of DSC in inset. **d, e**, Mean cumulative probability curves of DSC diameter (**g**) and length (**h**) compiled from the different imaging techniques; ExM (black, individual dendrites in grey), STED (blue) and EM (red) DSC properties were similar across modalities, ExM: DSCs diameters of 0.33±0.03 µm (range 0.11-0.46 µm) and length of 1.4±0.2 µm, n=35 DSCs in 6 dendrites; STED: microscopy DSC diameters of 0.36±0.01 µm (range 0.21-0.51 µm) and lengths of 2.8±0.3 µm, n=24 DSCs; EM reconstructions: DSC diameters of 0.26±0.06 µm and lengths of 1.6±0.7 µm; n=9 DSCs. f, **g**, Cumulative probability curves of DSC distance from soma and the dendritic sub-compartment size between two DSCs. Data determined from individual dendrites imaged using expansion microscopy (ExM; mean in black and each individual dendrite in grey). **h-k**, Dendritic shaft constrictions (DSCs) are present in cortical pyramidal neurons. **h**, Example of an entire reconstructed layer 2/3 pyramidal neuron, with the locations of the DSCs depicted in the insets indicated with number. Note the prevalence of DSCs in the apical tuft. Basal dendrites are shown in green, apical dendrites in blue, and DSCs indicated with red circles. **i, j**, Mean cumulative probability plots of DSC diameter (i) and length (j). **k**, Cumulative probability plots of DSC distance from soma.

To confirm the existence of DSCs in live tissue, we used STED microscopy and examined GC dendrites in mice expressing eYFP in granule cells (**Fig. 1b**, 8 slices from 3 animals). Here, we analyzed multiple dendritic segments from proximal (50-100 µm from the granule cell layer) and distal (>200 µm from the GC layer) dendritic regions (**Fig. 1b**). DSCs, defined as in the expansion microscopy experiments, were readily observed in the STED images (examples shown in **Fig. 1b**, right panels). As predicted by the distribution of DSCs in ExM experiments, DSCs occurred more frequently in distal (23/50) than proximal segments (1/15). Their dimensions were comparable to those measured by ExM (see **Fig. 1d, e**).

We used electron microscopy (EM) as a third method to detect DSCs in GCs (**Fig. 1c**). EM images were taken at distal locations (175±24 µm from the soma), and eighteen dendritic segments were reconstructed (mean length of segments 15±0.3 µm, range 6.3-31 µm). Nine DSCs were identified using the same detection criterion used for ExM and STED analyses (in 9/18 distal dendritic segments, n=2 mice, **Fig. 1c** lower panels, see **Extended Data Fig. 1f, g** for more examples). DSC dimensions obtained by EM were comparable to values from light microscopy (see **Fig. 1d, e**). In addition, EM images revealed subcellular features of the dendrites, such as the frequent presence of tubular structures within DSCs (**Fig. 1c, Extended Data Fig. 1g**), and the presence of mitochondria in adjacent dendritic shafts (**Extended Data Fig. 2**).

We next investigated if these structures are unique to dentate GCs. We first examined CA1 pyramidal neurons labelled using viral transduction with EGFP and anatomically reconstructed following ExM (**Extended Data Fig. 3a, b**). No constrictions were observed in the CA1 trunks up to 300 µm from the soma (n=2, n=1 mouse), however, DSCs were also observed in the fine radial oblique dendrites (n=3; see **Extended Data Fig. 3b, c**, DSC diameter and length in **Extended Data Fig. 3d**).

We then asked if DSCs also occur in cortical neurons. To this end, we examined serial EM from the MICrONS Consortium ^9^ dataset which contains complete reconstructions of entire cortical pyramidal neurons from layer 2/3 of the mouse visual cortex (**Fig. 1h**). Similar to CA1 pyramidal neurons, DSCs were restricted to the fine caliber dendrites with a range of 11-16 DSC in each cell (n=5 pyramidal neurons, n=1 animal; **Extended Data Fig. 4**). DSCs were primarily found in the apical dendritic compartment, with a majority occurring in the distal tufts (DSC locations indicated in a representative neuron in **Fig. 1h**). In total, 68 DSCs were identified, with 58 in apical and 10 in basal dendrites from 5 cells (corresponding to ∼85% of DSCs within a cell located in the apical dendrites, **Fig. 1h**). The average diameter at the DSC was 169.3±121.3 nm, ranging between 55 and 476 nm (**Fig. 1i**), and the average length of the constricted region was 1.2±0.4 µm, ranging between 0.3 and 1.8 µm (**Fig. 1j**). In keeping with the preferential localization in apical tufts, most DSCs were located distally, at an average distance of 190.3±87.8 µm from the soma, ranging between 13 and 360 µm (**Fig. 1k**).

Finally, we asked if DSCs occur across species. In human GCs labelled with GFP in organotypic slices (^10^, **Extended Data Fig. 1h**), we could also identify DSCs using identical criteria to the analyses in mice (DSC average diameter 0.57±0.05 µm, average length 2.3±0.9 µm, n=5 DSCs in 3 dendrites, **Extended Data Fig. 1i, j**).

In summary, using three different high-resolution microscopy approaches, we report on a novel morphological feature of fine caliber dendritic shafts in several different types of excitatory neurons, in mouse and human brain tissue.

### Computational modeling predicts altered dendritic integration of distal dendritic inputs

To predict the impact of DSCs on dendritic input integration, we generated a simplified ball and stick *in-silico* model. Two synaptic input sites were placed, one proximal to where DSCs occur, and one near the dendritic tip (90 and 220 µm from soma, respectively, **Fig. 2a**). Both sites featured mixed AMPA and NMDA conductances with a 1:1 NMDA/AMPA ratio. Local EPSPs were elicited at each input site (**Fig. 2a**). We then incorporated 1–7 DSCs into the model, matching their experimentally observed dimensions (200 nm width, 1 µm length, spaced 10 µm apart). This configuration introduced an axial resistance of 58–406 MΩ, consistent with values derived from our morphological data.

**Fig. 2:**
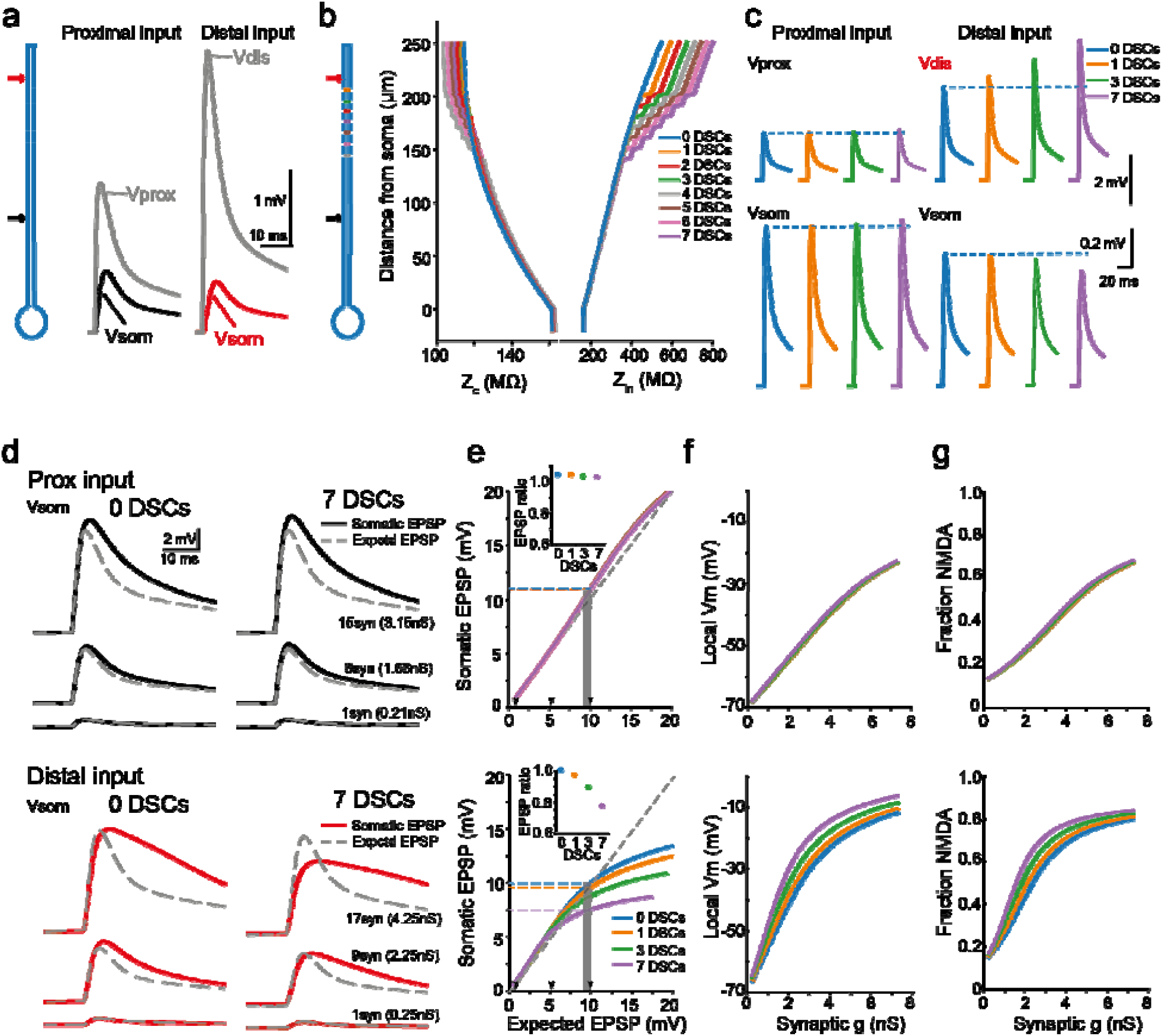
Computational modeling predicts modified dendritic integration of distal inputs. **a**, *left*, Schematic representation of ‘ball & stick’ model consisting of a soma (diameter 20 µm) and dendrite (length 250 µm diameter 1 µm), and showing position of proximal (black) and distal (red) input. *Right*, voltage traces measured at the indicated location in response to proximal or distal input showing attenuation of dendritic EPSP at the soma (synaptic conductance gMax = 0.26 nS for both, 90 and 220 µm from the soma, respectively). **b**, *left*, schematic representation of model showing position of 1-7 dendritic shaft constrictions (DSCs, length 1 µm and diameter 200 nm). *Right*, plots showing transfer impedance (Z_c_) and input impedance (Z_in_) at indicated position from soma with 0 to 7 DSCs. Z_in_ increased by 10 to 49% for 1 and 7 DSCs, respectively. **c**, voltage responses measured at indicated location in response to proximal or distal input with 0, 1, 3, or 7 DSCs as indicated. **d**, EPSPs measured at the soma in response to increasing input at proximal or distal inputs. Single synaptic conductance was 0.21 and 0.26 nS for proximal and distal sites and set as the conductance to elicit unitary EPSP amplitudes determined experimentally in **Extended Data Fig. 6**. Increasing the synaptic input was achieved by increasing the synaptic conductance in multiples of the single synaptic conductance. Number of model synapses and total conductance indicated. The expected EPSP, shown in grey, was determined by scaling the unitary EPSP by the number of model synaptic inputs. **e**, plots of somatic EPSP amplitude against expected EPSP amplitude for proximal (top) and distal (bottom) input with increasing number of DSCs from 0-7 (lines overlapping with proximal input). Dashed grey lines indicate unity gain. Arrowheads on x-axis correspond to traces shown in d. *Insets*, mean ratio of somatic EPSP amplitude to expected EPSP amplitude at an expected EPSP amplitude of 8-10 mV. **f**, plot depicting peak voltage and **g** fraction NMDA receptor recruitment against total synaptic conductance at the proximal (*top*) and distal input (*bottom*) sites with increasing number of DSCs (lines are overlapping with proximal input).

The addition of DSCs increased input impedance (Z_in_) and decreased transfer impedance (Z_c_) for distal input sites (**Fig. 2b**). Consequently, distal synaptic inputs exhibited enhanced local depolarization, with local EPSP amplitudes increase by 10% to 49%, depending on the number of DSCs added (**Fig. 2c**). Further, adding DSCs led to attenuation of distally evoked EPSPs measured at the soma by 1.4% to 14% with 1-7 DSCs, respectively (**Fig. 2c**, right traces). Proximal synaptic inputs, in contrast, were minimally affected (**Fig. 2c**, left traces). Lastly, we analyzed the scenario where synaptic inputs were confined to dendritic subsegments bordered by DSCs on both sides (see **Extended Data Fig. 5a, b**).

We next examined the impact of DSCs on synaptic summation, focusing on somatic voltages up to ∼20 mV (see **Extended Data Fig. 5a, e**). For proximal inputs, synaptic summation resulted in largely linear increases of somatic EPSP amplitudes (**Fig. 2d, e**, *upper panel*s). The presence of DSCs distal to the proximal synaptic input had little effect on this linear relationship even for large synaptic inputs (**Fig. 2d, e**, upper panel & inset).

In contrast, synaptic summation at distal input sites became markedly sublinear as the number of DSCs increased (**Fig. 2d, e**, lower panels). The EPSP ratio decreased from 1.01 without DSCs to 0.77 with 7 DSCs (**Fig. 2e**, lower panel inset). Notably, increasing the unitary synaptic conductance did not alter synaptic gain, as the relationship between measured and expected EPSPs remained consistent (**Extended Data Fig 5c**). The observed sublinear integration can be attributed to voltage saturation at distal input sites, driven by a high input impedance Z_in_ introduced by DSCs (schematic in **Extended Data Fig. 5f**). DSCs increased both distal dendritic Z_in_, and the local membrane depolarization across a range of stimulation strengths, to the point where voltage saturation was observed (**Fig. 3f**). These large local voltage changes led to a rapid recruitment of NMDA receptors in the presence of DSCs (**Fig. 2g**, lower panel).

**Fig. 3:**
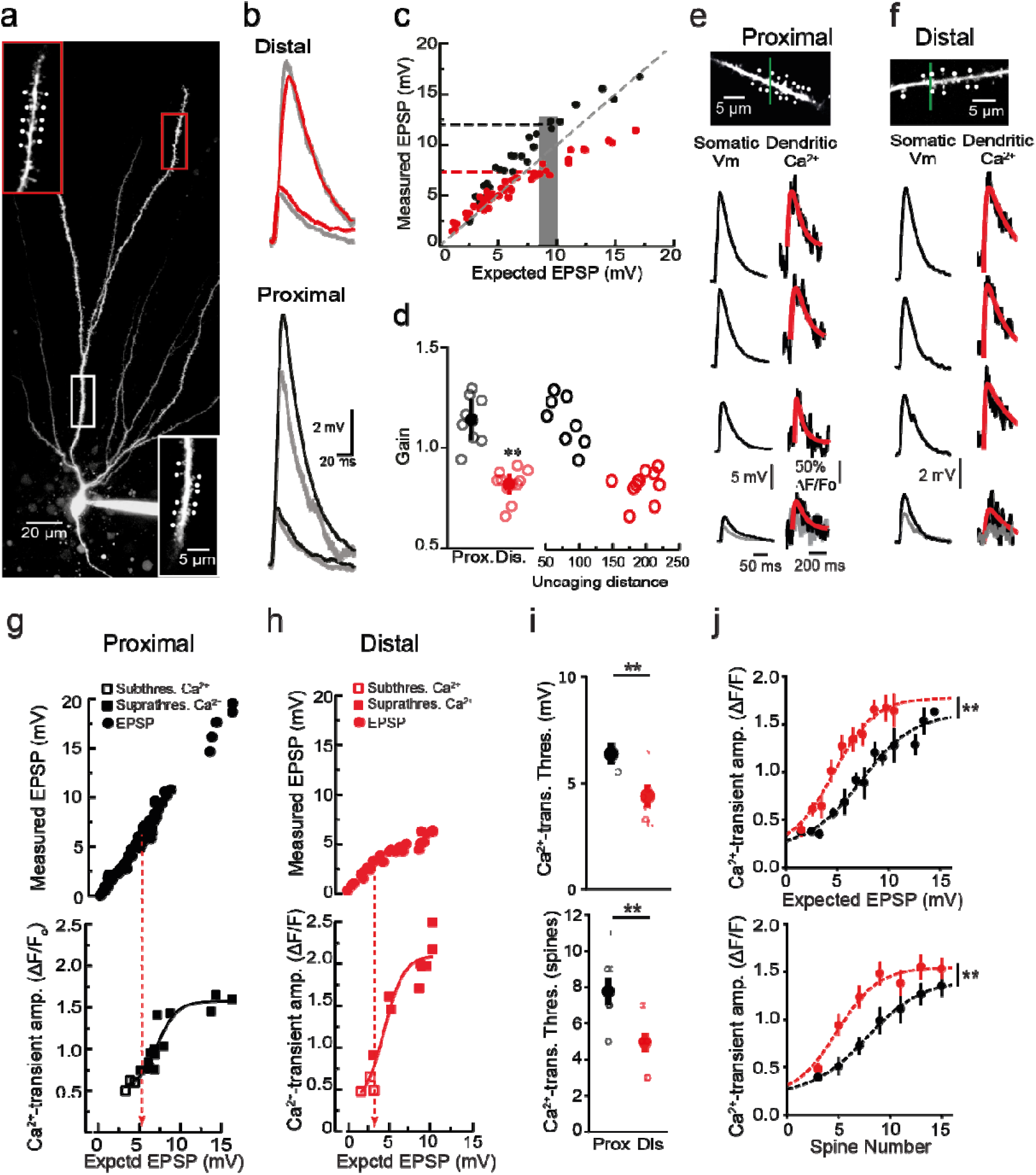
Input site specific integration of synchronous synaptic input. **a**, Granule cell filled with Alexa 594. Two-photon glutamate uncaging was performed at proximal (white box) and distal (red box) input sites. *Insets*: higher magnification images with uncaging points marked by dots. **b**, compound EPSPs evoked by near-synchronous glutamate uncaging of 2 and 10 spines at distal (red) and proximal (black) sites. In grey, expected compound EPSPs determined by arithmetically summing individual single spine responses. **c**, Plots of measured vs expected EPSP amplitudes for proximal (black) and distal (red) inputs for data shown in (b). Dashed grey line indicates unity gain. Grey bar indicates range of expected EPSPs used to compute gain, determined from mean ratio of measured vs. expected EPSP amplitudes over 8-10mV range (grey box). Gain of distal input (red dashed line) was lower than gain of proximal inputs (black dashed line) in same cell. **d**, summary plot of gain at proximal and distal input sites and with uncaging distance from some. **e, f**, Two-photon images of granule cell dendrites with uncaging sites (white spots) and line-scan location (green). Somatic EPSPs and the corresponding local dendritic Ca^2+^ linescans in response to increasing synaptic input at proximal or distal sites. Traces with suprathreshold (black) and subthrehold Ca^2+^ transients (grey). Suprathreshold Ca^2+^ transients were defined by a peak amplitude of >5x the baseline standard deviation and fit with an exponential rise and decay function (red). **g, h**, Plots of measured EPSP amplitudes (circles) and suprathreshold Ca^2+^ transient amplitudes (closed squares) against expected EPSP amplitudes. Subthreshold Ca^2+^ amplitudes shown in open squares, threshold indicated with dashed line. Ca^2+^ responses at proximal and distal input sites fit to a sigmoid function. **i**, Mean Ca^2+^ transient threshold measured as expected EPSP amplitude (mV) or spine number. Ca^2+^ threshold was significantly lower at distal vs proximal dendrites. Statistical significance determined using unpaired Student’s t-test distal vs. proximal sites, p<0.01 for both panels. **j**, Summary plot showing Ca^2+^-transient amplitude against expected EPSP amplitude or spine number. Mean Ca^2+^-transient amplitudes with sem binned in 1 mV bins of expected EPSP amplitude or spine number. Ca^2+^-transient amplitude were fit to sigmoid fits. Statistical significant difference determined between proximal and distal site using two-way ANOVA. Main effect proximal (n=4) vs. distal (n=9): F_(1,4)_ = 10.9, p<0.01; spine number F_(1,4)_=15.6, p<0.01; interaction: F_(1,4)_=1.2, p=0.34 and proximal (n=4) vs. distal (n=9): F_(1,9)_ = 26.8, p<0.01; expected EPSP F_(1,9)_=15.9, p<0.01; interaction: F_(1,9)_=0.4, p=0.93).

### Two-photon glutamate uncaging reveals sublinear summation, and enhanced NMDA receptor recruitment at distal GC dendrites

To experimentally test the predictions of the computational model, we used granule cells and performed two-photon glutamate uncaging at proximal and distal sites (78±17 vs. 195±14 µm from the soma, n=8 and 11, respectively, **Fig. 3a, b**, uEPSP properties in **Extended Data Fig. 6**). We found that input summation at proximal sites was linear with a gain above unity, as previously described (**Fig. 3c, d**, 1.14±0.10, n=8 dendrites, ^11^). In contrast, in keeping with the model predictions, distal site inputs were integrated sublinearly (0.82±0.05, n=11, p<0.01, Mann-Whitney U-test compared to distal sites). Sublinear integration at distal sites was not caused by dendritic expression of K^+^ channels (see **Extended Data Fig. 7a-c**).

We then assessed how steeply intradendritic Ca^2+^ transients depended on input strength at proximal and distal sites (**Fig. 3e, f**). We found that thresholds to generate Ca^2+^ transients were significantly lower in distal compared to proximal dendrites (**Fig. 3g, h, I**, n=13 and 16 for proximal and distal sites, respectively; both *p*<0.01 Student’s t-test), consistent with the modeling results. Additionally, Ca^2+^ transients reached saturation with fewer inputs and smaller expected EPSP amplitudes at distal input sites (two-way ANOVA p<0.01 for both, **Fig. 3g, h, j**). Blocking NMDARs with D-APV significantly increased Ca^2+^ transient thresholds (**Extended Data Fig. 8a, b**, Student’s t-test p<0.01) and shifted to the right the relationship between input and Ca^2+^ transient amplitudes (**Extended Data Fig. 8c**, two-way ANOVA p<0.01). These data show that distal dendrites exhibit strong sublinear summation and NMDA receptor recruitment, most likely due to local voltage saturation, as predicted by the computational model.

In summary, we identify dendritic shaft constrictions (DSCs) as a previously unrecognized morphological feature of dendrites. DSCs exert a strong influence on the passive electronic properties of dendrites, and shape dendritic integration. DSCs are found in different excitatory neuron types and exhibit selective dendritic distribution patterns. The selective occurrence of DSCs, combined with their consistent detection using three distinct methodological approaches, underscores their biological relevance rather than being artefactual. The molecular underpinnings of DSCs are unknown, but strong candidates are actin-dependent mechanisms relying on periodic cytoskeletal structures present in most dendrites ^12–14^. Alternatively, cell-cell interactions with surrounding neural elements could be responsible. The discovery of DSCs raises intriguing questions, such as whether DSCs are developmentally regulated, modulated by neuronal activity, or affect diffusion and active transport within dendritic shafts, akin to spine neck functions ^15,16^. Additionally, by enhancing NMDA-dependent Ca^2+^ signalling in distal dendrites, DSCs may shape plasticity rules unique to specific dendritic compartments ^17^. Our findings establish a foundation for exploring the roles of DSCs in dendritic function and synaptic plasticity.

## METHODS

### Animals

All experiments followed institutional guidelines of the Animal Care & Use committee of the University of Bonn. All in-vitro electrophysiological experiments were performed on adult (8-12 wks) C57BL/6J male wild-type mice (Charles River, Sulzfeld). For anatomical reconstructions of granule cells, C57BL/6J male wild-type mice were used for electron microscopy and expansion microscopy and (Thy1-YFP-H mice; Jackson labs strain 003782) were used for Stimulated Emission Depletion (STED) microscopy. For reconstructions of CA1 pyramidal neurons, we used of wt mice (C57BL/6 mouse line). Mice were kept under a light schedule of 12h on/12h off, constant temperature of 22±2°C, and humidity of 65%. They had ad libitum access to water and standard laboratory food at all times. All efforts were made to minimize animal suffering and to reduce the number of animals used.

### Viral injections

To sparsely label dentate granule cells, we injected retroviral constructs derived from a Moloney murine leukemia virus (MoMLV)-based vector with GFP expression driven by the chicken beta-actin (CAG) promoter. Wild-type animal (8-12 wk old) were anesthetized using a mixture of Fentanyl (0.05 mg/kg, Janssen-Cilag GmbH, Neuss, Germany), Midazolam (5 mg/kg, Ratiopharm GmbH, Ulm, Germany), and Medetomidin (0.5 mg/kg, Cp-Pharma, Burgdorf, Germany) and placed in a stereotactic apparatus (Neurostar, Tübingen, Germany). Small craniotomies were performed bilaterally at specific coordinates. About 1 µL of retroviral solution was injected (0.25 µL/min) into the dorsal dentate gyri via a Hamilton 1701 RN Syringe equipped with a Neuros Adapter and a blunt ended 30-gauge needle (Hamilton, Reno, NE). The following coordinates relative to Bregma were used: caudal 2 mm, lateral 1.5 mm, and ventral 2.1 mm. At the end of the procedure the anaesthesia was antagonized by a mixture of Naloxon (1.2 mg/kg, Ratiopharm GmbH), Flumazenil (0.5 mg/kg, Hameln Pharma plus GmbH, Hameln, Germany), and Atipamezol (2.5 mg/kg, Orion Pharma GmbH, Hamburg, Germany). Carprofen (5 mg/kg, Pfizer GmbH, Berlin, Germany) was administered as postoperative analgesic and Enrofloxacin (5mg/kg, Bayer Vital GmbH, Leverkusen, Germany) as antibiotic. Mice were sacrificed for immunohistochemistry ∼6wk after injection to allow for maturation of granule cells.

To sparsely label CA1 pyramidal neurons, we infected the hippocampal CA1 region of wt mice (C57BL/6 mouse line) with a rAAV-expressing EGFP under control of the human synapsin1 promoter (rAAV-syn-GFP) by stereotaxic virus injection (WPI Benchmark/Kopf; coordinates: rostro caudal: 2.1□□mm from the Bregma; mediolateral: ±1.2□□mm from the midline; dorsoventral: 2.1□□mm) and stained afterward with an antibody against EGFP. The use of low virus titers (∼10^8^/ml) resulted in a stochastic and relatively sparse labelling of CA1 pyramidal neurons.

### Expansion Microscopy

The expansion microscopy protocol was adopted from ^18,19^. In short, the labelled hippocampal slices up to 200 µm thick were incubated with 2 mM methylacrylic acid-NHS linker for 24 hours on a shaker at room temperature (RT). After washing three times in PBS, the slices were incubated overnight (ON) in the monomer solution (8.6% sodium acrylate, 2.5% acrylamide, 0.15% N,N’-methylene bisacrylamide, and 11.7% NaCl in 1× PBS) on a shaker at 4°C. The gelling solution was prepared by adding 4-hydroxy-TEMPO (0.01%), TEMED (0.2%) and ammonium persulfate (0.2%) to fresh monomer solution. During gelling, the hippocampal slices were placed in a 24-well plate on ice to avoid early polymerization. After applying the gelling solution, samples were put on a shaker at 4°C for 5 minutes and then be transferred to the gelling chamber, followed by 3 hours incubation at 37°C. After the gel formation, the samples were incubated at 37°C in the digestion buffer (50 mM Tris, 1 mM EDTA, 0.5% Triton-X100, 0.8 M guanidine HCl, and 16 U⁄mL of proteinase K; pH 8.0), exchanging the buffer every 24 hours if necessary. In general, a 100 µm thick sample takes about 16 h to be completely digested. After digestion, the buffer was removed, and the samples were washed three times with PBS. The preservation of the fluorescence of autofluorescent proteins during the expansion procedure enabled us to perform immunolabeling of the GFP-expressing cells after clearing the samples with the digestion.

Immunostaining for GFP was carried out as follows. To prevent unspecific binding of the primary antibody, the samples were incubated with blocking buffer (1x PBS, 5% normal goat serum, 0.3% TritonX-100, 0.02% sodium azide) on a shaker ON at RT. After blocking, the samples were incubated for 24 h in blocking buffer with the primary antibody (chicken anti-GFP; 1:500 in blocking buffer; Abcam, ab13970) on a shaker at 4°C. The following day, slices were washed at RT in blocking buffer three times for 30 min and incubated 24 h in secondary antibody (goat anti-chicken conjugated with Alexa Fluor® 488, 1:400 in blocking buffer; Thermo Scientific, A-11039) on a shaker at 4°C. For nuclear staining, all samples were stained using Hoechst 33342 (H3570, Invitrogen).

Expanded and stained samples were imaged on a custom light-sheet microscope. In short, for fluorescence excitation four fiber-coupled lasers emitting at 405, 488, 561 and 638 nm (Hübner Photonics, Germany) were employed. The horizontally scanned light sheet was generated by a galvanometer system with silver-coated mirrors. The adjustment of the beam waist position within the sample chamber was realized by a relay optics mounted on a linear precision stage. The beam waist in the object plane was adjusted to a 1/e^2^ diameter of 6.5±0.02 µm for the 405 nm, 7.3±0.02 µm for the 488 nm, 7.0 ± 0.02 µm for the 561 nm, and 8.3±0.02 µm for the 638 nm laser lines. For illumination we used a Mitutoyo 10x NA 0.28 air objective. Our custom-designed sample chamber featured an illumination window formed by a conventional 24×24 mm coverslip with a thickness of 0.17 mm. The sample was observed from the top using a Nikon 25x NA 1.1 water immersion objective (CFI75 Apochromat 25XC W) with an additional 1.5x magnification (Nikon, Germany). The sample was mounted on a coverslip, which could be moved in three spatial directions by motorized micro-translation stages. We used a sCMOS camera (3200 × 3200 pixels, pixel size 6.5 μm, Kinetix, Teledyne Photometrics, USA) for data acquisition in global shutter mode yielding a pixel size (x, y) of 0.173 µm and a step size (z) of 0.3 in expanded tissue. For analysis, pixel sizes were corrected for the expansion factor of the individual slice (see analysis below). All electronic components were controlled by a custom-written LabView program.

The sample size greatly exceeded the visual field size of 554 μm^2^. Therefore, the data were acquired in a mosaic-like fashion, ensuring a 15% spatial overlap between two neighboring tiles. Complete 3D representations of the samples were possible after several 3D data sets were stitched together using Fiji ^20^ and the stitching plugin of ^21^. In order to optimize the stitching process, especially when datasets exceed the available RAM memory of the workstation, the process was performed in two steps. First, substacks of the 3-D data sets were created using a Fiji script. Each substack contained about 15% of the information located in the center of the full stack. Secondly, each substack was stitched to its respective neighboring substack yielding the best overlap in terms of the cross-correlation measure. Based on the localization information of each substack after stitching, the full 3D stacks were stitched.

Selected image stacks were spatially deconvolved using Huygens (Professional version 22.04, Scientific Volume Imaging, The Netherlands). Deconvolution was performed using theoretical PSFs, based on microscopic parameters. The classical maximum likelihood estimation algorithm was used with Acuity: 0.0, and a SNR value between 8.0 to 20.0 for a maximum number of iterations between 75 to 100 were selected. Stitching and deconvolution was performed on a workstation equipped with two Intel Xeon Gold 6252 CPU (2.1 GHz, 24 cores), 1 TB memory, and an Nvidia RTX A4000 GPU (16 GB GDDR6) running under Windows Server 2019.

### Preparation of acute brain slices

Transgenic animals (Thy1-YFP-H line), 21–40 days old, were killed by cervical dislocation, and their brains were quickly removed and placed in ice-cold sucrose-based artificial cerebrospinal fluid (ACSF) containing (in mM) 210 sucrose, 10 glucose, 2 KCl, 26 NaHCO_3_, 1.25 NaH_2_PO_4_, 0.1 CaCl_2_, and 6 MgCl_2_ (pH 7.4, osmolarity ∼320 mOsm/L), which was bubbled with carbogen (95% O_2_/5% CO_2_). Sagittal 350-μm-thick hippocampal slices were cut using a vibratome (VT1200, Leica, Mannheim, Germany) and transferred to a heated (32°C) holding chamber with NaCl-based ACSF bubbled with carbogen, which consisted of (in mM) 124 NaCl, 3 KCl, 26 NaHCO_3_, 1.25 NaH_2_PO_4_, 10 glucose, 2 CaCl_2_, 1 MgCl_2_, and 0.6

Trolox (pH 7.4, osmolarity ∼305 mOsm/L) for 1 h. Slices were subsequently maintained at room temperature for a maximum of 4 h. For the imaging experiments, slices were transferred to a submerged recording chamber, where they were continuously perfused (2.1 mL/min) with ACSF at room temperature.

### Stimulated Emission Depletion (STED) Microscopy

We imaged the acute hippocampal slices using a home-built STED microscope based on 2-photon excitation and stimulated emission depletion (STED) using pulsed lasers ^22,23^. Briefly, a femtosecond mode-locked Ti:Sapphire laser (Chameleon, Coherent, Santa Clara, CA) operating at 80 MHz and emitting light at 834 nm was used in combination with an optical parametric oscillator (OPO BASIC Ring fs, APE, Berlin, Germany) to produce STED light pulses at 598 nm that were stretched out to ∼100 ps using a 20 m-long polarization-maintaining fiber. A STED light reflection served to synchronize a second Ti:Sapphire laser (Tsunami, Spectra Physics, Darmstadt, Germany) tuned to 910 nm and running at a repetition rate of 80 MHz, which was used for 2-photon excitation of the fluorophores. The STED doughnut was created by passing the STED beam through a vortex phase mask (RPC Photonics, Rochester, NY), which imposed a helical 2π-phase delay on the wave front. Wave plates (λ/2 and λ/4) were used to make the STED light circularly polarized before it entered into the objective. The 2P and STED beam were combined using a long-pass dichroic mirror. The two laser beams were moved over the sample in all three dimensions using a x-y galvo scanner (Janus IV, TILL Photonics) and a z-axis nano-positioner (Pifoc, PI, Karlsruhe, Germany). A water-immersion objective with a long working distance (1.5 mm) equipped with a correction collar was used (60X LUMFI, 1.1 NA, Olympus). The correction collar was adjusted to optimize the STED doughnut using gold nanospheres (diameter=150 nm; BBInternational, Cardiff, United Kingdom) and slightly readjusted as a function of imaging depth. The epi-fluorescence was de-scanned and imaged onto an avalanche photodiode (SPCM-AQR-13-FC, PerkinElmer, Villebone-sur-Yvette, France). Signal detection and peripheral hardware were controlled by the Imspector scanning software (Abberior Instruments, Göttingen, Germany) via a data acquisition card (PCIe-6259, National Instruments). Regions of interest (20 × 20 μm^2^) were imaged with a pixel size of 10 nm and pixel dwell time of 50 μs. The laser power at the sample was typically around 5–20 mW for 2P and 5–15 mW for STED.

### Electron Microscopy

Mouse brain tissue was fixed with 4% paraformaldehyde and 2.5% glutaraldehyde in cacodylate buffer. Samples were initially incubated with reduced osmium (1% osmium, 0.8% potassium ferricyanide, cacodylate buffer) then washed and incubated in either 320mM pyrogallol in water or 1% thiocarbohydrazide in water. Afterwards samples went through a second non-reduce osmium staining (1% osmium in water). Then samples were incubated overnight in 1% uranyl acetate in water. Next day, they were incubated in Walton’s lead aspartate. After washing and dehydration samples were embedded in Epon or Durcupan resin. Serial ultrathin sections (50 nm) were cut on a RMC Powertome and collected on carbon coated glass slides or formvar carbon coated copper slot TEM grids. Imaging of the samples was done in a Zeiss Crossbeam 550 with backscattered electron detector or STEM detector, respectively. Consecutive sections were scanned and registered with ATLAS 5 software to gain a z stack of the same consecutive regions. Each image was 5001 × 5001 pixels with an x-y pixel size of 10 nm. The serial images were rigidly registered using the Fiji plugin Linear Stack Alignment with SIFT (https://imagej.net/plugins/linear-stack-alignment-with-sift) and then processed using IMOD version 4.11.20 ^24^ to perform segmentation, isosurface rendering and 3D reconstructions.

#### Morphological analysis of cortical pyramidal neurons reconstructed by serial electron microscopy

The neuron IDs used in this study were downloaded from the *MICrONS* dataset ^9^ and include: 864691135693733567 (Figure 1h), 864691136194996812, 864691135374673737,864691135738411121, and 864691134884807418 (version 943). Neuronal skeletons were generated using *Ultraliser* ^*25*^ with a segment resolution of 75 nm. DSCs were automatically detected based on the following criteria: a local reduction in dendritic shaft diameter of at least twofold, followed by a recovery to at least 75% of the original diameter within a distance of no more than 3 µm. Additionally, the absolute reduction in diameter was required to exceed 500 nm. All detected DSCs were manually validated using the 3D dendritic reconstruction. The analysis code is available on GitHub.

#### Confocal microscopy of human organotypic hippocampal brain slice cultures

The preparation and cultivation of human organotypic hippocampal brain slice cultures were carried out according to the protocol described previously ^10^. At 1 day *in vitro* (DIV) the slices were transduced with 1 μl rAAV-hSyn-eGFP (Addgene #50465-AAVrg). At DIV15, the slices were fixed in 4% paraformaldehyde (PFA) and mounted in Fluoromount-G (SouthernBiotech). Confocal imaging was performed using an inverted Zeiss LSM 980 confocal laser scanning microscope with Airyscan 2, along with the Zeiss ZEN blue 3.6 software with the ZEN Module Airyscan Joint Deconvolution. Overview images of the dentate gyrus were taken using the Plan-Apochromat M27 10x (N.A. 0.45) objective with a laser wavelength of 488 nm. For detailed imaging, regions with a high fluorescence level in sparsely populated areas of the medial-distal part of the apical dendrite were selected. Z-stacks were captured with a laser wavelength of 488 nm using the Plan-Apochromat 40x (N.A. 1.3) oil immersion objective with a 4x zoom and a step size of 0.18 μm. The images underwent further processing using the Airyscan processing and joint deconvolution with 10 iterations, as provided by Zeiss ZEN blue 3.6 software.

### Analysis of DSC morphology

Individual dendrites were binarized and dendritic diameters determined in Fiji using the Local Thickness plugin. Local Thickness was extracted at the center of the dendrite using a skeleton created from the binarized image using the Fiji plugin Skeletonize (2D/3D) ^26^. Fluorescent images of dendrites were filtered to reduce noise using the 3d median filter plugin ^27^ with a kernel radii of 1,1,0.2 in pixels. The images were binarized using a threshold automatically set using the MaxEntropy method in Fiji and adjusted manually if necessary. Open holes closed using 2-3 iterations of Dilate followed by Fill Holes and the Erode functions. Following binarization, all binary images were superimposed with original grey scale images, overlap manually compared and deviations corrected. Binary images were scaled such that the x,y and z scales were identical using the scale plugin in Fiji and missing places interpolated using the Bicubic average process. Using the Local Thickness plugin, binarized images were filled with the largest non-overlapping spheres (**Extended Data Fig. 1b**). The binarized image was also smoothed with 3D gaussian filters and skeletonized to provide a path through the middle of the dendrite that was used to extract the diameter of the largest sphere at each location along the dendrite using MATLAB. Data was then analysed in python. The extracted diameters were smooth using a boxcar filter over 10 pixels. Dendritic shaft constrictions were identified as drops in diameter that were a factor of 2. Edges of the constrictions were where the sharp changes in 1^st^ derivative of the diameter returned to values below 20nm/pixel. The diameter had to recover by at least 75% of the original value. In the case of expansion microscopy, diameters were corrected for expansion. The expansion factor experimentally determined for each tissue section by comparing somatic and dendritic images obtained prior to and following expansion (see **Extended Data Fig. 1a**). Pre-expansion images were enlarged on average±SD by 3.95±0.16 times [range: 3.8-4.1] to optimally overlap with post-expansion images.

### Electrophysiology - Slice preparation and patch-clamp recording

Mice were deeply anesthetized with isoflurane and then decapitated. Brains were rapidly removed and placed in ice cold (<2°C) sucrose-based artificial cerebrospinal fluid (sucrose-ACSF) containing (in mM): 60 NaCl, 100 sucrose, 2.5 KCl, 1.25 NaH_2_PO_4_, 26 NaHCO_3_, 1 CaCl_2_, 5 MgCl_2_, 20 glucose. Horizontal slices of 300 μm were cut with a vibratome (Leica VT1200S, Wetzlar, Germany) and incubated in sucrose-ACSF at 35°C for 30 min. Subsequently, slices were transferred to a submerged holding chamber containing normal ACSF containing (in mM): 125 NaCl, 3.5 KCl, 1.25 NaH_2_PO_4_, 26 NaHCO_3_, 2 CaCl_2_, 2 MgCl_2_, 15 glucose at room temperature. All extracellular solutions were equilibrated with 95% O_2_ and 5% CO_2_.

Cells within the suprapyramidal layer were visualised with infrared oblique illumination optics and a water immersion objective (60x, 0.9 NA, Olympus). All cells were clearly in the granule cell layer and we targeted cells in the first third of the granule cell layer that displayed a more oblong shape to record from cells with apical and basal processes in plane with the slice surface to ensure dendrites were not cut. Somatic whole-cell current-clamp recordings were performed with a BVC-700 amplifier (Dagan Corporation, Minneapolis MN, USA). Data were filtered at 10 kHz and sampled at 50 kHz with a Digidata 1440 interface controlled by pClamp Software (Molecular Devices, Union City, CA). Patch-pipettes were pulled from brosilicate glass (outer diameter 1.5 mm, inner diameter 0.86 mm; Science Products, Hofheim, Germany) with a Flaming/Brown P-97 Puller (Sutter Instruments, Novato, USA) to resistances of 2 to 5 MΩ in bath and series resistances ranging from 8 to 30 MΩ. Only cells with an input resistance <400 MΩ were selected to exclude recordings from newly generated granule cells ^28^. The standard internal solution contained (in mM): 140 K-gluconate, 7 KCl, 5 HEPES, 0.5 MgCl_2_, 5 phosphocreatine, 0.16 EGTA. Internal solutions were titrated to pH 7.3 with KOH, had an osmolality of 295 mOsm, and contained 50-100 μM Alexa Fluor 594 (Invitrogen, Eugene OR, USA). In experiments to record synaptic-induced intracellular Ca^2+^ changes in the dendrite, Cal-520 (200 µM; Invitrogen) replaced EGTA and recordings were begun following 20 min, to allow equilibration of the dye in the cell. Voltages were not corrected for the calculated liquid junction potential of +14.5 mV. Membrane potential was adjusted to -70 mV for all recordings. Active and passive properties were determined from hyperpolarizing and depolarizing current steps (800 ms) of increasing amplitudes injected via the somatic patch pipette. Passive membrane properties, action potential properties and firing patterns were assessed throughout the entire course of the experiment. Cells with unstable input resistances or lacking overshooting action potentials were discarded as well as recordings with holding currents >-200 pA for 70 mV and access resistances > 30 MOhm.

### Two-photon uncaging

Two-photon glutamate uncaging at apical dendrites of dentate granule cells was performed using a dual galvanometer-based scanning system (Prairie Technologies, Middleton, WI, USA) to photo-release glutamate at multiple dendritic spines. MNI-caged-L-glutamate 15 mM (Tocris Cookson) was dissolved in HEPES-buffered solution (in mM as follows: 140 NaCl, 3KCl, 2 MgCl_2_, 2 CaCl_2_ 20 D-glucose, and 10 HEPES, pH 7.4 adjusted with NaOH, 305 mOsmol/kg) and was applied using positive pressure via glass pipettes (< 1 MΩ) placed in closed proximity to the selected apical dendrite. Distance measurements were performed on the maximum projection of image stacks collected at the end of recordings using Fiji (ImageJ, NIH). The distance between the soma and the input site was measured as the length of line from the centre of the soma along the dendrite to the approximate midpoint of the input site. All uncaging locations included in this study were located >25 µm <200 µm from the soma. We used two ultrafast laser beams of Ti:sapphire pulsed lasers (Chameleon Ultra, Coherent). One pulsed laser tuned to 820 nm to excite the Alexa 594 and select 10–15 dendritic spines in close vicinity (∼10 μm in length); another tuned to 730 nm to photo-release MNI-glutamate at the preselected spines. The intensity of each laser beam was independently controlled with electro-optical modulators (Conoptics Model 302RM, Danbury, CT, USA). MNI-glutamate was uncaged at increasing number of spines (2-15) with a 1 ms exposure times and the laser was rapidly moved from spine to spine with a transit time of ∼0.1 ms. In any group of spines, uncaging was always performed from the most distal toward the most proximal spine. The laser power at the slice surface was kept below 22 mW to avoid photo damage. The midpoint of the stimulated dendritic region was assessed using maximum projection images from two-photon stacks, and the 2D distance determined.

The glutamate was uncaged onto a sequence of single spines to elicit uncaging evoked unitary excitatory post synaptic potentials (uEPSPs). To quantify deviations from linearity in dendritic integration, the arithmetic summation calculated from each individual single spine uEPSP was compared to the actually measured compound EPSP during glutamate uncaging onto the same sequence of spines.

### Ca^2+^ imaging

The calcium dye Cal-520 (200 µM) was loaded into the cell via the patch-pipette and the cell filled for at least 20 mins after break-in before recording. Cal-520 was used for all experiments as we found Cal-520 had higher signal to noise compared with or OGB-1 (200 µM; see **Extended Data Fig. 9**). Two-photon imaging of Cal-520 and Alexa594 (50 µM) was performed using a pulsed laser tuned to 820 nm, the fluorescent emissions collected via a water-immersion objective (60x, 0.9 NA, Olympus) and signals separated by a dichroic mirror (DXC 575). Red Alexa-594 emission collected at 605/45, and green Ca^2+^ dye emission collected at 525/70 were detected using separate photomultiplier tubes (Hamamatsu). With these filters, red Alexa-594 emission was not detected above noise in the green channel. Simultaneous whole-cell somatic recording and local dendritic calcium imaging (linescans at 425–750 Hz) were conducted. For action potential induced Ca^2+^ transients, APs were triggered by a 1-5 short current injections (2 ms) in a train (at 100Hz) through the patch pipette. Each trial was repeated at least 3 times and the mean value is presented. For uncaging evoked EPSP induced Ca^2+^ transients, the line-scan was centrally placed between spine number 3 and 5. To reduce photodamage during simultaneous two-photon Ca^2+^ imaging and uncaging, line-scans were performed in response to uncaging on 3,5,7,9,10,12,14,15 spines. In addition, we used a passive pulse-splitter to convert each uncaging laser pulse into 8 sub-pulses to further reduce photodamage ^29^. For iontophoresis induced Ca^2+^ transients the line-scan was placed across the proximal dendrite near the iontophoretic pipette. Intra-dendritic Ca^2+^ transients are reported as changes in Cal-520 fluorescence (ΔF/F_o_), normalised to baseline fluorescence (F_o_) averaged over the 190 ms period prior to stimulation. Calcium transients were distinguished from noise as a fluorescence peak at least 4x above the standard deviation of the baseline period and were fit by a double exponential with a rise tau within the range of 1-50 ms and a decay tau within the range of 80-500 ms. The reported Ca^2+^ transients increase linearly over a ΔF/F_o_ range of 1 to 5 (n=16, see **Extended Data Fig. 9**) suggesting the Cal-520 dye fluorescent changes did not saturate over this range (also see ^30^).

### Modeling

A simple multicompartmental passive ‘ball and stick’ model with number of segments following the d_lambda rule ^31^ and passive properties Ra = 181 Ωcm, Cm = 1 uFcm^−2^ and a leak conductance = 0.0002 Scm^−2^, which gave an Rin of 165 MΩ, were adopted from Carnevale and Hines (2010) ^31^ and Krueppel et al. ^10^(2011). The model is available on GitHub (https://github.com/tonykelly00/DSC_model). A soma (20 µm diameter) contained one or two dendrites (1 µm diameter, 250 µm length). The transfer (Zc) and input impedance (Zn) were determined from the model. Simulations were run in the Neuron 7.8 simulation environment.

Synaptic conductances were modelled using point mechanisms for AMPA and NMDA conductacnces and were placed on the dendrite at 90 and 220 µm from the soma. The AMPA conductance was modelled with a biexponential function (AMPA tau_rise = 0.05 ms, AMPA tau_decay = 2 ms) and the NMDA conductance modelled with a biexponential function with a voltage-dependent Mg^2+^ block (NMDA tau_rise = 0.33 ms and NMDA tau_decay = 50 ms). The Mg^2+^ block of NMDA conductance was depended on voltage with a sensitivity of gamma = 0.06 mV^-1^ and a on Mg^2+^ concentration with a sensitivity of eta = 0.05 mM^-1^.

The range of unitary synaptic conductances (0.1–1 nS; see ^32^) elicited EPSP amplitudes in the model, which covered the range of somatic EPSP amplitudes that were experimentally measured (0.2 mV – 2.0 mV, see **Extended Data Fig. 6**). A unitary synaptic conductance was selected that elicited a unitary EPSP equivalent to the average unitary EPSP measured experimentally at the soma. Multiples of the unitary EPSPs conductance were used to simulate compound EPSPs (measured EPSP) and the amplitude compared to the arithmetic sum of the unitary response (expected EPSP). It should be noted that the relationship between measured and expected EPSP amplitudes was insensitive to the unitary synaptic conductance (gSyn; **Extended Data Fig. 5c**) over the unitary somatic EPSP amplitudes typically observed experimentally (unitary EPSP amplitudes of 0.2-2 mV see **Extended Data Fig. 6**). To assess the synaptic gain we determined the EPSP ratio as the measured EPSP amplitude relative to the expected EPSP over the range of expected EPSPs amplitudes of 8-10 mV. In experiments assessing the interaction of proximal and distal inputs, NMDA recruitment fraction is the fraction of total NMDA receptors recruited. Dendritic shaft constrictions were modelled as short (1 µm) dendritic segments with a diameter of 0.2 µm and 1-7 DSCs were located between 140-200µm from the soma at 10 µm intervals.

## Data and code availability

Data used to generate the figures and examples of raw data is available at Zenodo (https://zenodo.org/records/14037025?preview=1&token=eyJhbGciOiJIUzUxMiJ9.eyJpZCI6ImY4NDE2YWU2LTlmMmEtNGY4NS04M2EwLTg4NzhhMTY1NGRhYSIsImRhdGEiOnt9LCJyYW5kb20iOiJiMGI0YmUwYTg3MjI3MThiNjU5MmY4YjVkOGE3MDE5MiJ9.3r6XoHH6eZQwoqN2nKAh0WgprCR1NjNeP7sbtyx3r9XMjD7o_HtmfZ2qxccQYmhidxqBO5on_0IdoWTX1FDYQg). Code used to analyse the data is also available in the Zenodo repository as well as GitHub (https://github.com/tonykelly00/DSC_morphology_pipeline).

## Acknowledgements

Supported by the research group FOR-2751 to TK and HB from the Deutsche Forschungsgemeinschaft (DFG), SFB 1089, Project C04 to HB and TT from the DFG, and SPP 2041 Computational Connectomics of the DFG to HB, MS and UK. The Joachim Herz Foundation funded CWC. We would like to thank Hannes Beckert and Lydia Maus at the Microscopy Core Facility of the Medical Faculty at the University of Bonn for providing support and instrumentation funded by the Deutsche Forschungsgemeinschaft (DFG, German Research Foundation) – Projektnummer 388171357. We thank Lea Keller for excellent technical assistance, Morton Fejer, Lasse Stephan Keuneke, Michelle Latham for help with analysis of morphological data and Isabelle Blameuser for help with EM reconstructions.

