## Supplemental Figures for "Nanoscale dendritic shaft constrictions shape synaptic integration in fine caliber principal neuron dendrites"

Extended Data Figures

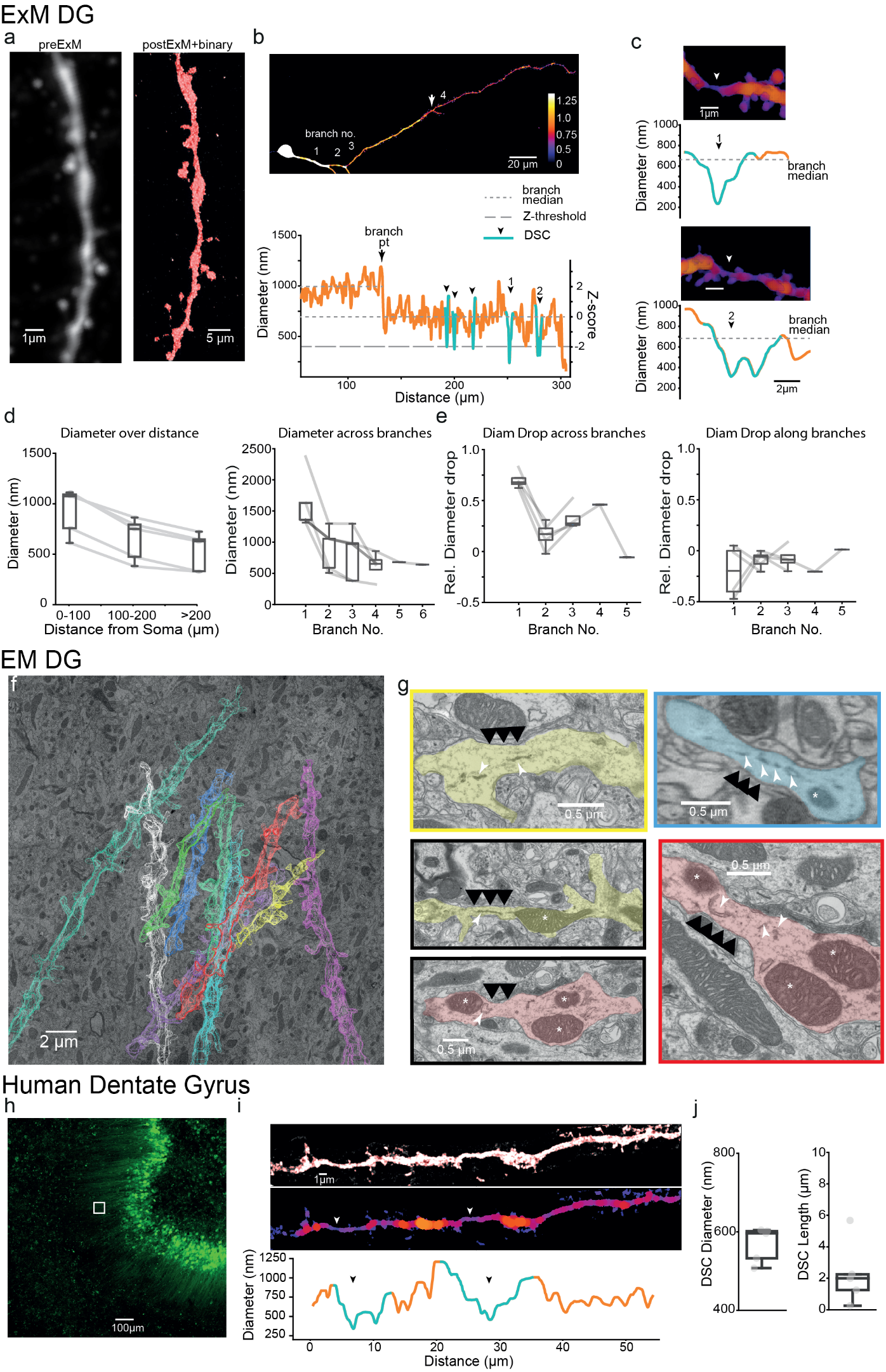

**Extended Data Fig. 1:** **Imaging granule cell dendrites at nano-scale resolution. a**, left: confocal maximum projection image of granule cell dendritic segment prior to expansion (preExM); right: maximum projection image with overlapping binary image (red) of same segment following ExM. Comparing preExM and postExM images was used to determine the expansion factor of each section. **b**, Example of a 3D reconstruction of an entire individual dendrite from soma to distal tip. Granule cell was sparsely labelled with eGFP using Molony virus transduction. Pseudo-colored with the diameter of the largest non-overlapping sphere (*above*) and plot of diameter against distance from soma for the distal half of the dendrite containing DSCs (*below*). At branch points (arrow) diameters drop as in this example, and remain at similar levels more distally. Drops in diameter by factor of >2 which re-increase to at least 75% were identified as DSCs indicated with cyan lines and arrowheads. Scale is corrected for using the expansion factor. Z-score of diameter shown on right axis. The z-score was calculated from the distribution of diameters obtained from the last dendritic branch. **c**, Example DSCs from b pseudo-colored with the diameter. DSCs indicated with arrows. Scale bar is corrected for the expansion factor. **d,** plots of average shaft diameter binned by distance from soma or branch No. **e**, Plots showing drop in diameter across branches determined as mean diameter at last 10 µm of parent branch compared to beginning 10 µm of daughter branch and drop in diameter along a branch determined as mean diameter at beginning 10 µm compared to last 10 µm. Drops in diameter expressed as proportions of total drop in diameter over entire dendrite. **f**, Transmission electron microscope image with overlay of segmented dendrites taken from serial imaging of 50 nm thick sections. **g**, example electron micrographs from 2 different stacks of the outer molecular layer (193 and 160 µm from the granule cell layer). Black arrowheads indicate DSCs where the diameter dropped by >2 compared to the neighboring dendritic shaft. White arrowheads indicate smooth tubular structures and white asterisks indicate mitochondria. **h**, confocal maximum projection image of human hippocampal organotypic slice (15 DIV) with dentate granule cells viral transduced with GFP. **i**, granule cell dendrite from g indicated with white box (155 µm from the granule cell layer) reconstructed, binarized and diameter plotted below. DSCs where diameter dropped by >2 indicated by white arrow and highlighted in cyan. **j**, Summary plots of DSC diameter and length for 5 identified DSCs

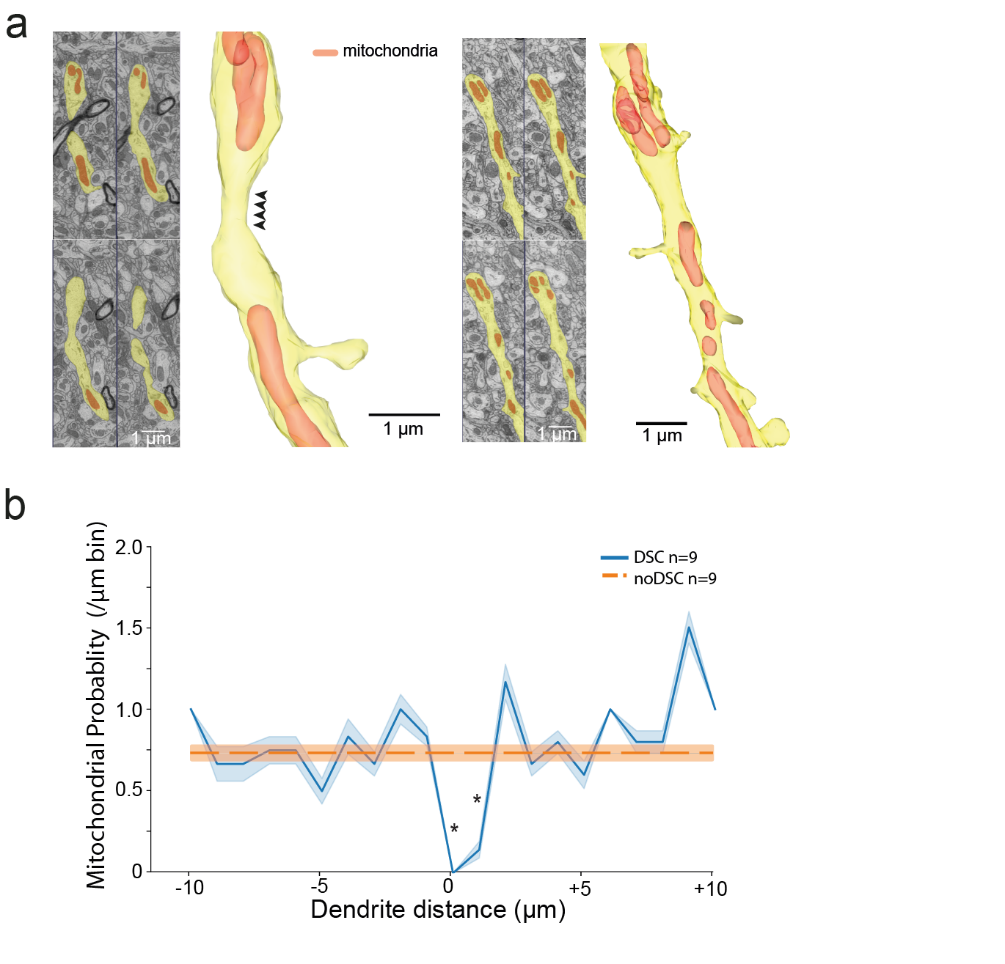

**Extended Data Fig. 2: Relation of DSCs to the presence of mitochondria. a**, *left*, example micrographs and 3d reconstructions of dendrites showing the absence of mitochondria at DSCs. DSC indicated with arrows. *Right*, also showing dendritic subsegments without mitochondria and without a DSC. **b**, the average mitochondrial density (mean ± sem) in 1 µm bins along dendritic segments containing DSCs, compared to the average mitochondrial density along the entire dendritic segment from all dendritic segments devoid of DSCs. Dendritic segments containing a DSC are aligned with the DSC at 0 µm. The mitochondrial probability was significantly reduced at the site of the DSC compared to the average mitochondrial density from all dendritic segments devoid of DSCs (*, indicates p<0.05, Dunnett’s test). However, the mere absence of mitochondria was not sufficient for a DSC, as most dendritic subsegments without mitochondria did not exhibit a DSC (14/23, length of 1.7±0.2 µm, n=14).

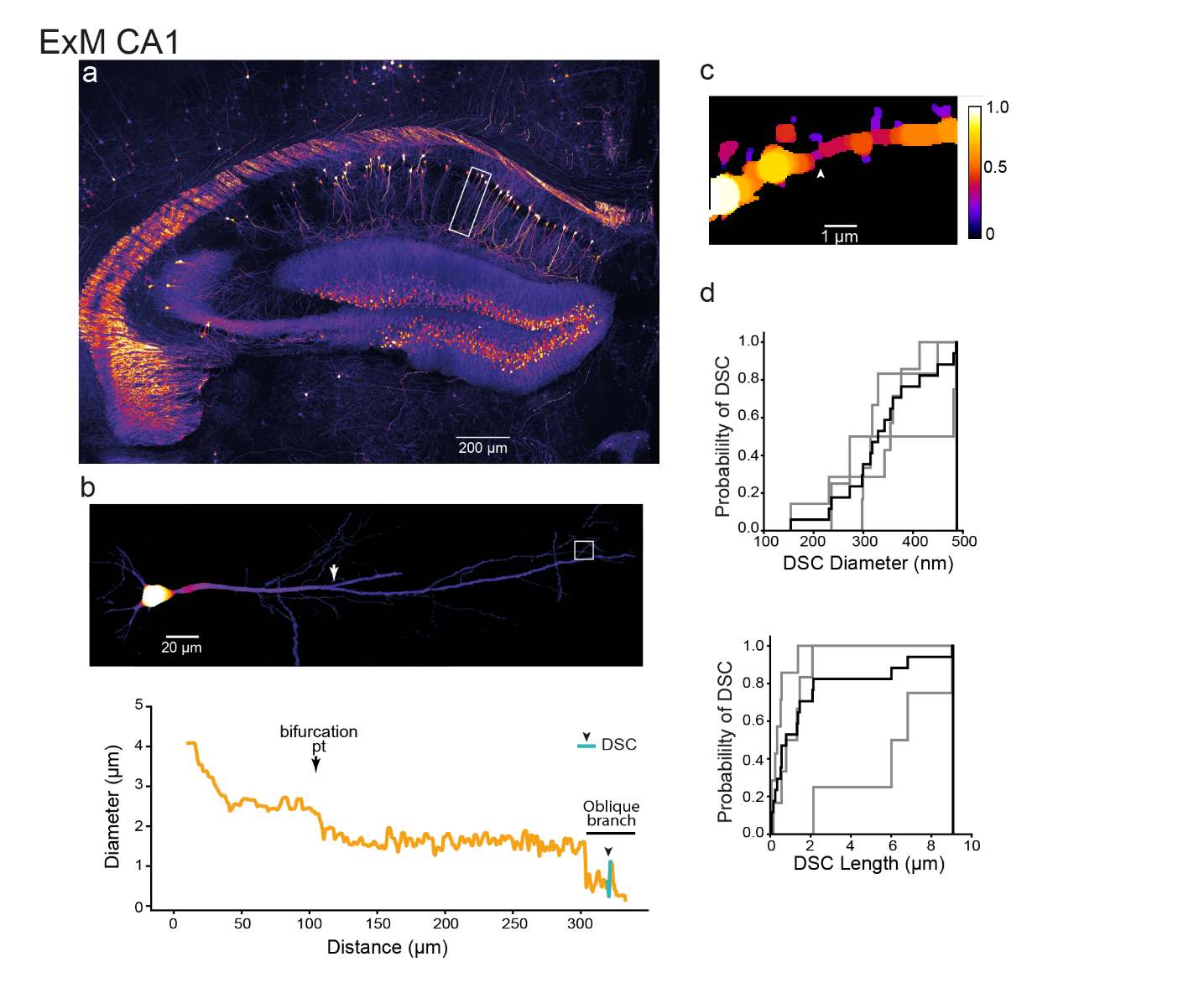

**Extended Data Figure 3: Expansion Microscopy of CA1 pyramidal cell dendrites. a**, Maximum projection image of expanded hippocampus acquired using light-sheet microscopy, scale bar corrected for expansion. White box outlines cell shown in b. **b**, Example of a 3d reconstruction of a CA1 pyramidal cell pseudo-colored image, white box indicates oblique branch (above), and plot of diameter against distance from soma (below). **c**, Example DSC from a CA1 pyramidal cell radial oblique dendrite. **d**, CA1 DSCs had an average diameter ± SD of 0.3±0.09 µm (range 0.15-0.48 µm, n=17 DSCs in 3 radial oblique dendrites) and an average length ± SD of 1.7±1.4 µm (range 0.5-9 µm, n=17 DSCs in 3 dendrites).

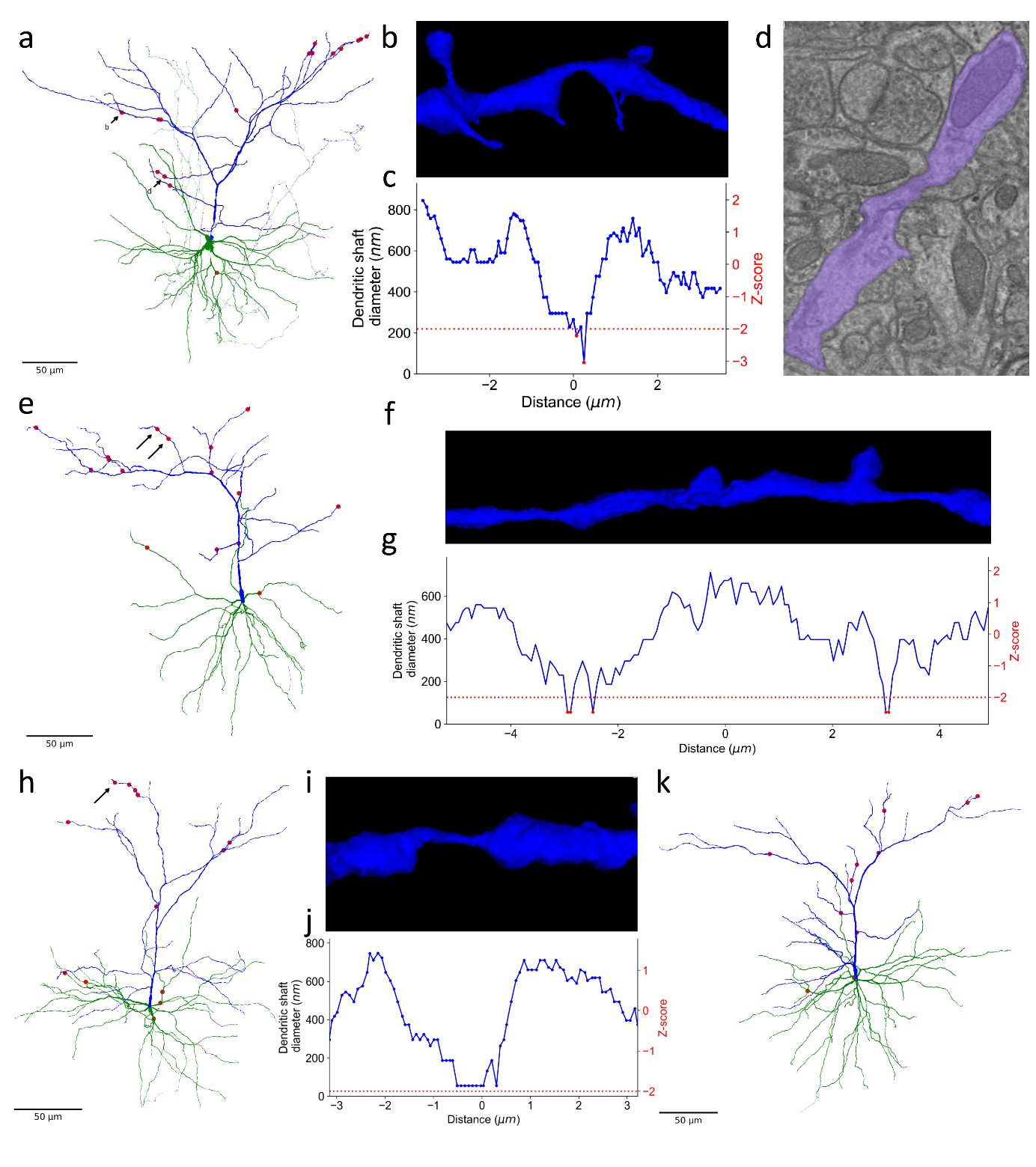

**Extended Data Figure 4: DSCs of cortical layer 2/3 pyramidal neurons from serial section EM a, e, h, k**, example reconstructions of pyramidal neurons with locations of DSCs indicated with red circles. **b, f, i,** example reconstructions showing DSCs, **d**, EM photomicrograph of DSC and **c, g, j** correspondine diameter plots with z score

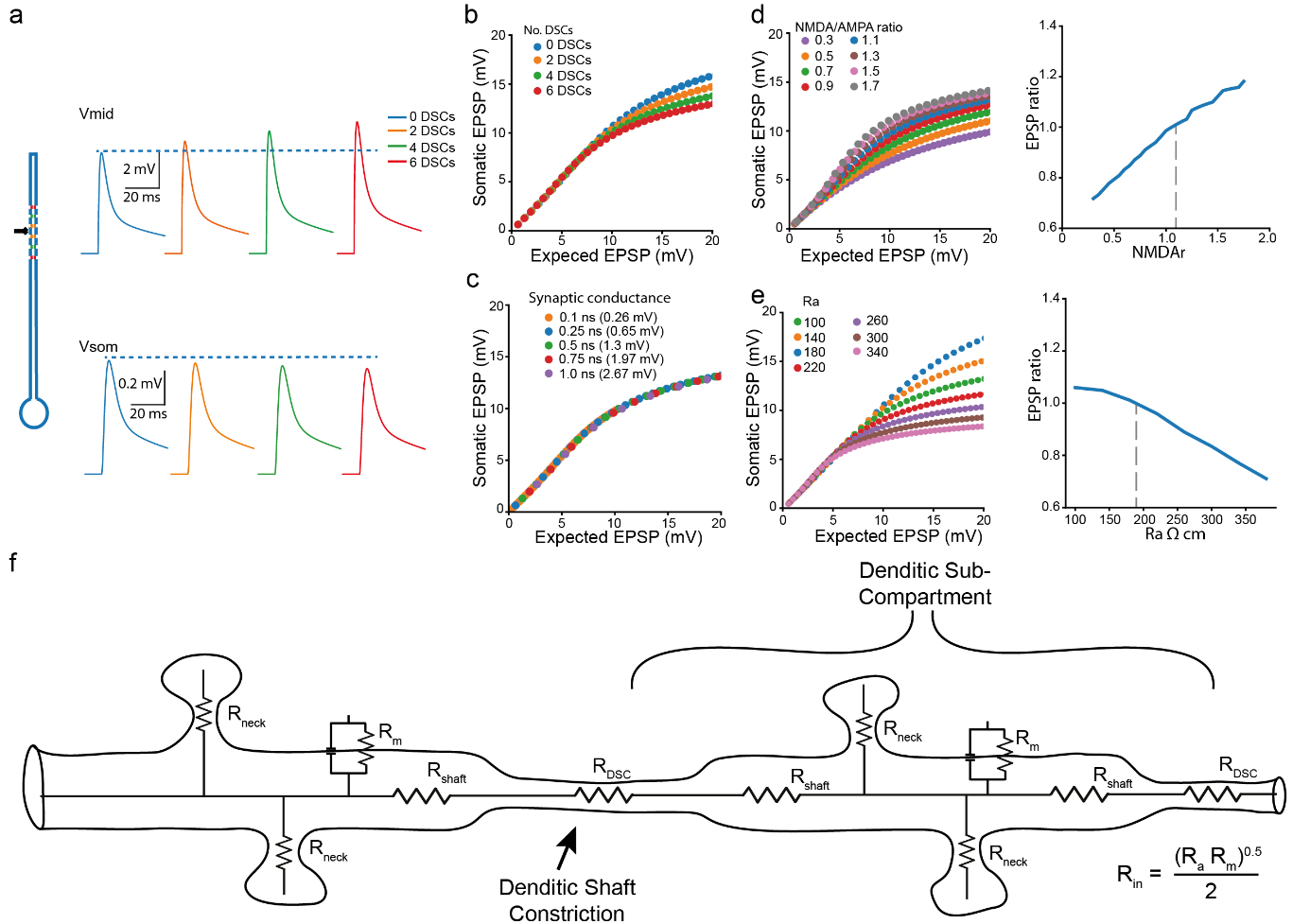

**Extended Data Figure 5: Effect of DSC position and neuronal parameters on dendritic integration.** **a**, *left*, Schematic representation of ‘ball & stick’ model consisting of a soma (diameter 20 µm) and dendrite (length 250 µm diameter 1 µm), and showing position of synaptic input (black; 165 µm from the soma) in the dendritic compartment and position of DSCs. *Right*, voltage traces measured at the indicated location in response to synaptic input (synaptic conductance g_Max_=0.26 nS). The local EPSP amplitude within the subsegment increased (amplitude increases by 8 to 28% when bordered on each side by 2 DSCs (1 on each side), 4 DSCs (2 each side), or 6 DSCs (3 each side), respectively and the efficiency of voltage propagation to the soma decreased (somatic EPSP amplitude decreased by 2 to 7% when bordered by on each side by 2 to 6 DSCs, respectively) **b**, Plots of somatic EPSP amplitude against calculated EPSP amplitude for synaptic input compartmentalised within DSCs (2,4,6 DSCs) or in the absence of DSCs. **c-e,** Effects of synaptic conductance (c), NMDA/AMPA ratio (d), and axial resistance (d) on the relationship between somatic EPSPs and expected EPSP amplitudes in the absence of any DSCs. **c,** Plots of somatic EPSP amplitude against expected EPSP amplitude showing that changing g_Syn_ does not change this relationship in the absence of DSCs. **d,** *Left panel:* Impact of large changes in NMDA/AMPA ratio (indicated in different colors, see legend) on the relationship of somatic EPSP amplitude to expected EPSP amplitude. *left panel*, a NMDA/AMPA ratio of 1.1 (see Krueppel et al. 2011) was decreased or increased and resulted in increases and decreases in synaptic gain. Synaptic gain determined as mean EPSP ratio of measured vs calculated EPSP amplitudes over the 8-10 mV range of expected EPSPs. *Right panel:* Plot of relationship between the NMDA/AMPA ratio and synaptic gain. Previously reported 1.1 value indicated by dashed line (Krueppel et al. 2011). **e,** *Left panel:* Impact of changes in axial resistance (R_a_) on relationship between somatic EPSP amplitude vs. expected EPSP amplitude. *Right panel*, plot of the relationship between R_a_ and synaptic gain. R_a_ of 180 Ω cm derived from literature indicated as dashed line (Krueppel et al. 2011, also see Stuart and Spruston 1998; Golding et al. 2005; Schmidt-Hieber et al. 2007). **f**, Schematic of dendritic segment with dendritic sub-compartment boarded by DSCs. The input impedance or electrical resistance of a compartment depends not only on the membrane resistance but also on the ease by which current flows into the compartment ie the input resistance (R_in_) of the sub-compartment is related to the membrane (R_m_) and axial resistances (R_shaft_ + R_DSC_) of the compartment. Reductions in diameter increase R_in_ due to increases in both R_m_ (Ω•cm) and R_a_ (Ω/cm). R_a_ is more dramatically increased due to the squared relationship of axial resistance to diameter (R_a_= 4R_A_/π•d^2^; R_m_ = R_M_/π•d; where R_M_ and R_A_ are the specific membrane (Ω•cm^2^) and axial (Ω•cm) resistances and d is diameter) (see Koch 1998; Segev 1998).

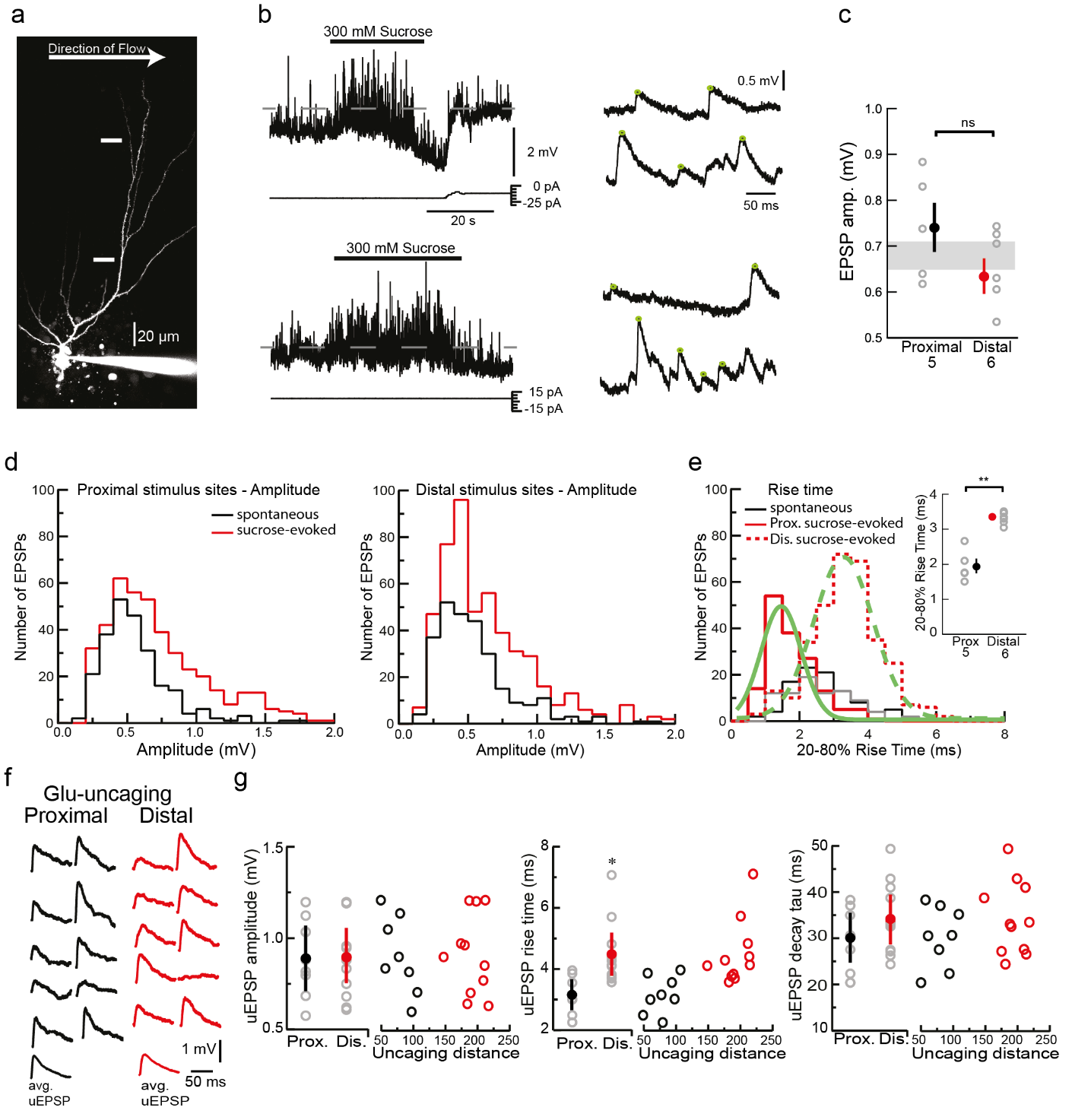

**Extended Data Fig. 6: Unitary EPSP properties elicited by sucrose or glutamate uncaging at proximal and distal input sites. a**, Two-photon maximum projection fluorescent image of granule cell filled with Alexa 594 via a patch-pipette. White lines indicate positions of sucrose pipette. **b**, Sucrose-evoked EPSPs at proximal and distal sites. **c** mean amplitudes of sucrose-evoked EPSPs were similar at proximal and distal sites (mean ± sem; Student’s t-test p=0.1). Grey shaded area indicates the mean(±sem) of 0.68±0.03 across all input sites and was used for computational model. **d**, distribution of sucrose-evoked EPSPs amplitudes at two inputs site. **e**, rise-time of sucrose-evoked EPSPs differed between proximal and distal input sites inset shows mean rise times for input sites (Student’s t-test p<0.01). **f**, single spine EPSPs evoked by glutamate uncaging at distal (red) and proximal (black) input sites. **g**, properties of individual single spine responses. The uEPSPs exhibited slower rise times at the distal site, as expected (3.2±0.5 ms vs. 4.5±0.7 ms, n=8 vs 11, for proximal and distal sites respectively; p<0.01 Student’s t-test). Between proximal and distal sites there was no significant difference in the uEPSP amplitudes (0.89±0.18 mV vs 0.90±0.15 mV, n=8 and 11; Student’s t-test p=0.94) or decay time constants (30.2±5.5 ms vs. 34±5.4 ms, n=8 vs 11, for proximal and distal sites respectively; p=0.22 Student’s t-test).

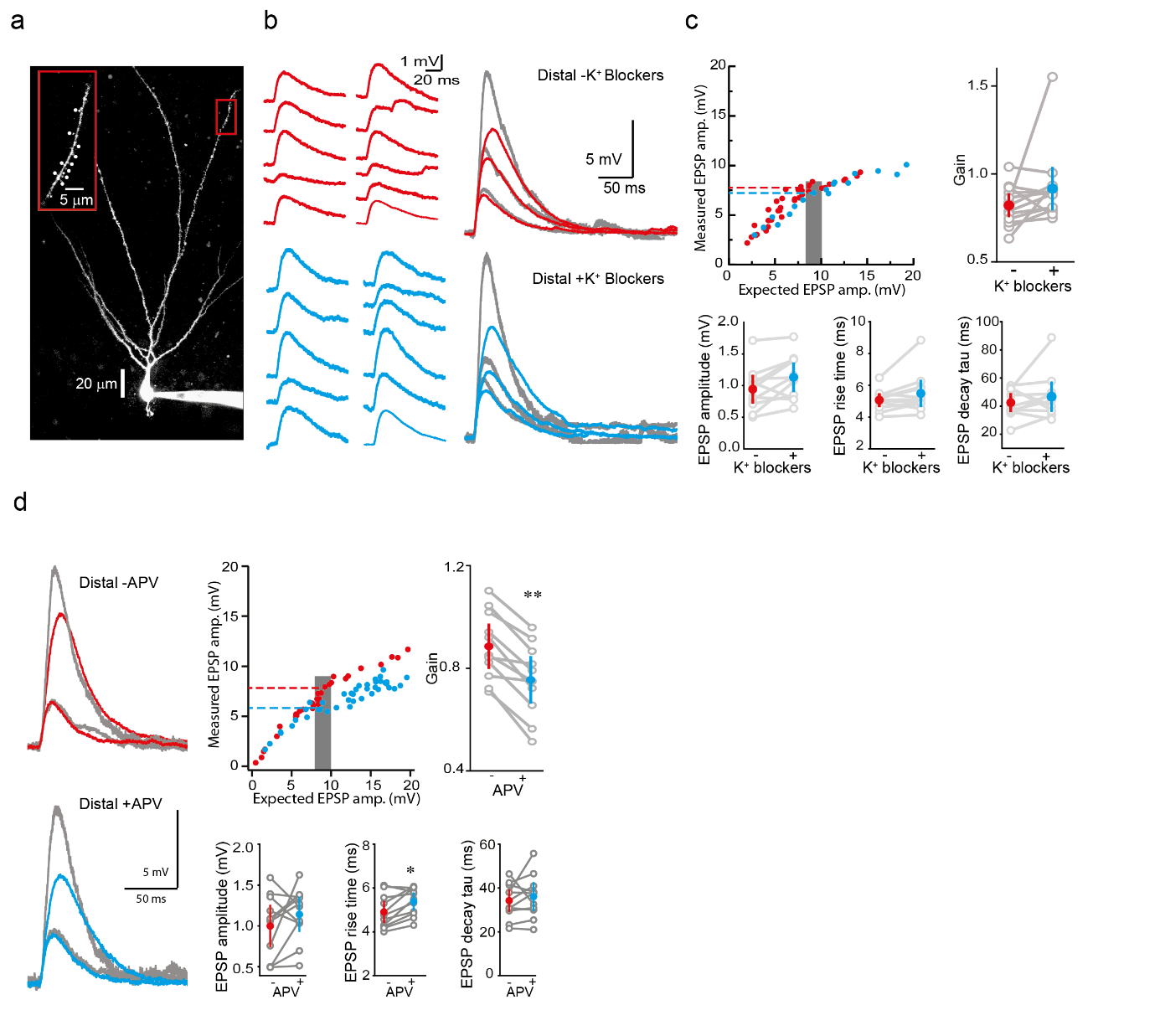

**Extended Data Fig. 7: Involvement of voltage-gated and synaptic conductances a**, Two photon image of granule cell filled with Alexa 594. Two-photon glutamate uncaging was performed at distal (red box) input sites. *Insets*: higher magnification images with uncaging points marked by dots. **b**, *left*, single spine EPSPs evoked by glutamate uncaging at distal input sites in the absence (red) and presence (blue) of K^+^ channel blockers (1 mM TEA and 200 µM 4-AP). *Right*, compound EPSPs evoked by near-synchronous glutamate uncaging of 2, 5 and 10 spines at distal sites. In grey, expected compound EPSPs determined by arithmetically summing individual single spine responses. **c**, measured versus expected EPSP amplitudes for data shown in (b). Grey bar indicates range of expected EPSPs used to compute gain determined from mean ratio of measured vs expected EPSP amplitudes over 8-10mV range (grey box). Gain of distal input was similar in the absence (red dashed line) and presence (blue dashed line) of K^+^ channel blockers (gain of 0.82±0.07 vs. 0.91±0.13 in the absence vs. presence of K^+^ channel blockers, n=12; p=0.18, paired Wilcoxon signed rank test). Properties of individual single spine responses.

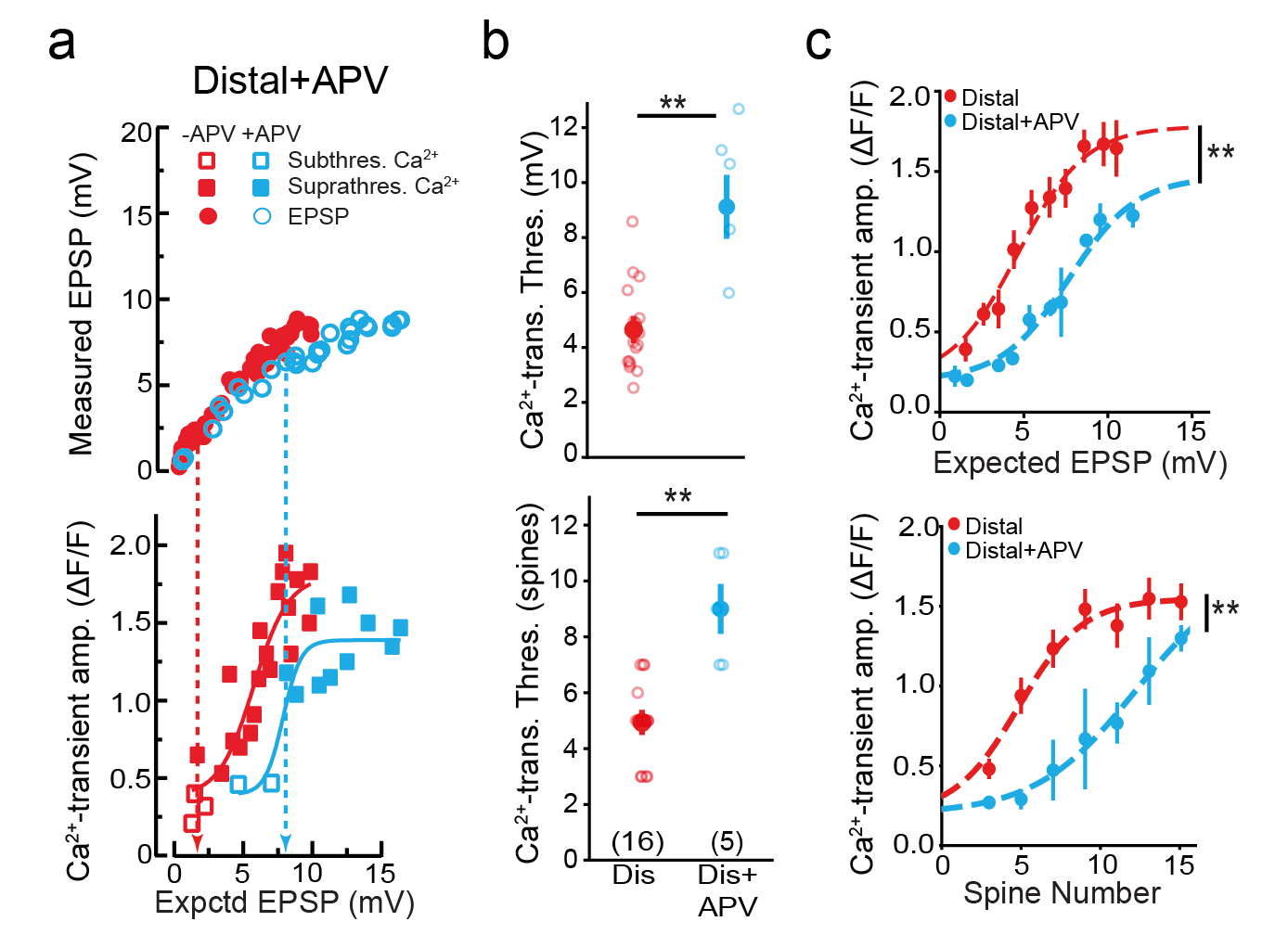

**Extended Data Fig. 8: Distal NMDA receptors mediate calcium influx into distal dendrites. a**, At distal sites, application of APV (blue) shifted threshold for suprathreshold Ca^2+^ responses to the right. **b,** Mean Ca^2+^ transient threshold measured as expected EPSP amplitude (mV) (from i) or spine number. Distal Ca^2+^ threshold significantly increased in the presence of APV. Statistical significance determined using one-way unpaired Student’s t-test distal sites vs. distal sites +APV, p<0.01 for both panels. **c**, Summary plot showing Ca^2+^-transient amplitude against expected EPSP amplitude or spine number. Mean Ca^2+^-transient amplitudes with sem binned in 1 mV bins of expected EPSP amplitude or spine number. Ca^2+^-transient amplitude were fit to sigmoid fits. Statistical significance determined using two-way ANOVA with Bonferroni’s post-test. Main effect distal (n=9) vs. distal+APV (n=3): F_(1,4)_ = 22.9, p<0.01; spine number F_(1,4)_=11.2, p<0.01; interaction: F_(1,4)_=0.8, p=0.51; two-way ANOVA main effect distal (n=9) vs. distal+APV (n=3): F_(1,8)_ = 34.7, p<0.01; expected EPSP, F_(1,8)_=18.5, p<0.01; interaction: F_(1,8)_=0.8, p=0.62).

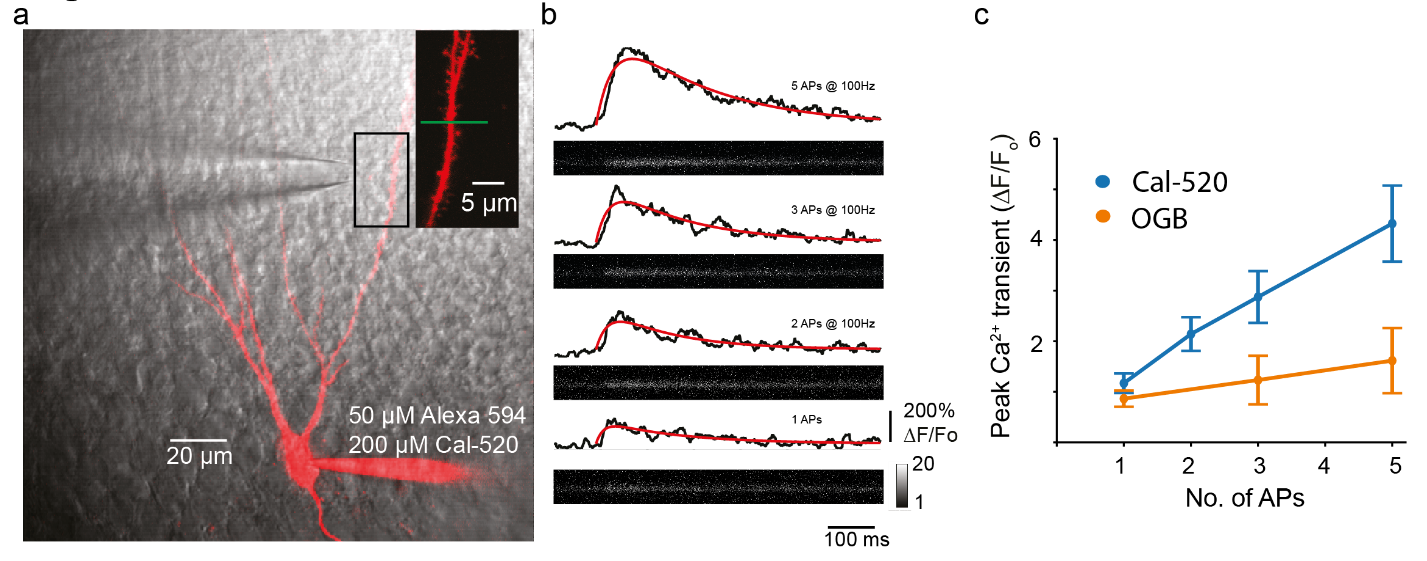

**Extended Data Fig. 9: Linear increases in Cal-520 fluorescence a**, Two-photon image of granule cell filled with Alexa 594 and calcium indicator Cal-520. Application pipette for MNI glutamate visible in transmitted channel. Intra-dendritic Ca^2+^ recorded at proximal dendrite. *Inset*: higher magnification image with line-scan position indicated (green line). **b**, Dendritic Ca^2+^ responses in response to somatically-evoked APs. **c**, Increasing the number of APs elicited linear increases in peak Ca^2+^ transient using both Cal-520 and OGB-1. Peak responses were significantly larger with Cal-520 and remained linear up to a DeltaF/F_o_ of at least 4.

Publication bibliography

Golding, Nace L.; Mickus, Timothy J.; Katz, Yael; Kath, William L.; Spruston, Nelson (2005): Factors mediating powerful voltage attenuation along CA1 pyramidal neuron dendrites. In *The Journal of physiology* 568 (Pt 1), pp. 69–82. DOI: 10.1113/jphysiol.2005.086793.

Koch, Christof (1998): Biophysics of Computation: Oxford University Press.

Krueppel, Roland; Remy, Stefan; Beck, Heinz (2011): Dendritic integration in hippocampal dentate granule cells. In *Neuron* 71 (3), pp. 512–528. DOI: 10.1016/j.neuron.2011.05.043.

Schmidt-Hieber, Christoph; Jonas, Peter; Bischofberger, Josef (2007): Subthreshold dendritic signal processing and coincidence detection in dentate gyrus granule cells. In *Journal of Neuroscience* 27 (31), pp. 8430–8441. DOI: 10.1523/JNEUROSCI.1787-07.2007.

Segev, Idan (1998): Cable and Compartmental Models of Dendritic Trees. In James M. Bower, David Beeman (Eds.): The Book of GENESIS. New York, NY: Springer New York, pp. 51–77.

Stuart, G.; Spruston, N. (1998): Determinants of voltage attenuation in neocortical pyramidal neuron dendrites. In *Journal of Neuroscience* 18 (10), pp. 3501–3510. DOI: 10.1523/JNEUROSCI.18-10-03501.1998.
